# An integrative cardio-informatics approach identifies positionally conserved ancestors of human lncRNA genes transcribed across vertebrate hearts

**DOI:** 10.64898/2026.10.21.759660

**Authors:** Anne-Marie Galow, Athanasia Pavlopoulou, Elif Yıldız, Stefan Kuhn, Stephan Marquardt, Joao Seco, Oleksandra Chabanovska, Praveen Vasudevan, Catharina Nesselmann, Christian Etz, Alexander Rebl, Robert David, Stella Logotheti

**Author notes:** These authors contributed equally (Anne-Marie Galow, Athanasia Pavlopoulou). Correspondence: Stella Logotheti; Robert David.

## Abstract

**Background:** The poor conservation of lncRNA genes limits their comprehensive functional studies, as genetic engineering of relevant animal models presupposes the presence of corresponding defined orthologs. This limitation is prominently evident in cardiovascular research, where heart-related lncRNAs are rarely conserved between humans and other vertebrates, despite the highly conserved nature of cardiac function. Because lncRNAs frequently regulate neighboring protein-coding genes essential for cardomyocyte function (herein termed cardiomyocyte indispensable, CMIs), we hypothesized that they are under evolutionary pressure to retain their genomic position and transcriptional activity, irrespective of sequence similarities to prototypical human lncRNA genes.

**Results:** By integrating mouse cardiac knockout phenotypes with human single-cell transcriptomics data, we systematically identified heart-expressed CMIs and their neighboring. lncRNA genes. Cross-species synteny analyses revealed regions flanking 109 CMIs with conserved relative positions and transcriptional orientations across vertebrate genomes. As most of these loci reside within poorly annotated non-coding regions, we developed *VertLncRNAchecker,* a neural network-based web interface, that predicts lncRNA-like genes within these regions, which we termed “positional ancestors”. Finally, we validated *in situ* that these positional ancestors express mature lncRNA transcripts in the hearts of mice, pigs, and frogs, mirroring the expression of their human lncRNA counterparts.

**Conclusions:** We present an accurate and retrainable computational platform for cross-species lncRNA discovery that prioritizes conserved genomic position over sequence conservation or existing genome annotations. This pipeline revealed putative ancestors of human heart-related lncRNAs hidden within the vertebrate non-coding “dark genome”, providing novel insights into lncRNA evolution and a useful resource for cardiovascular research.

GRAPHICAL ABSTRACT

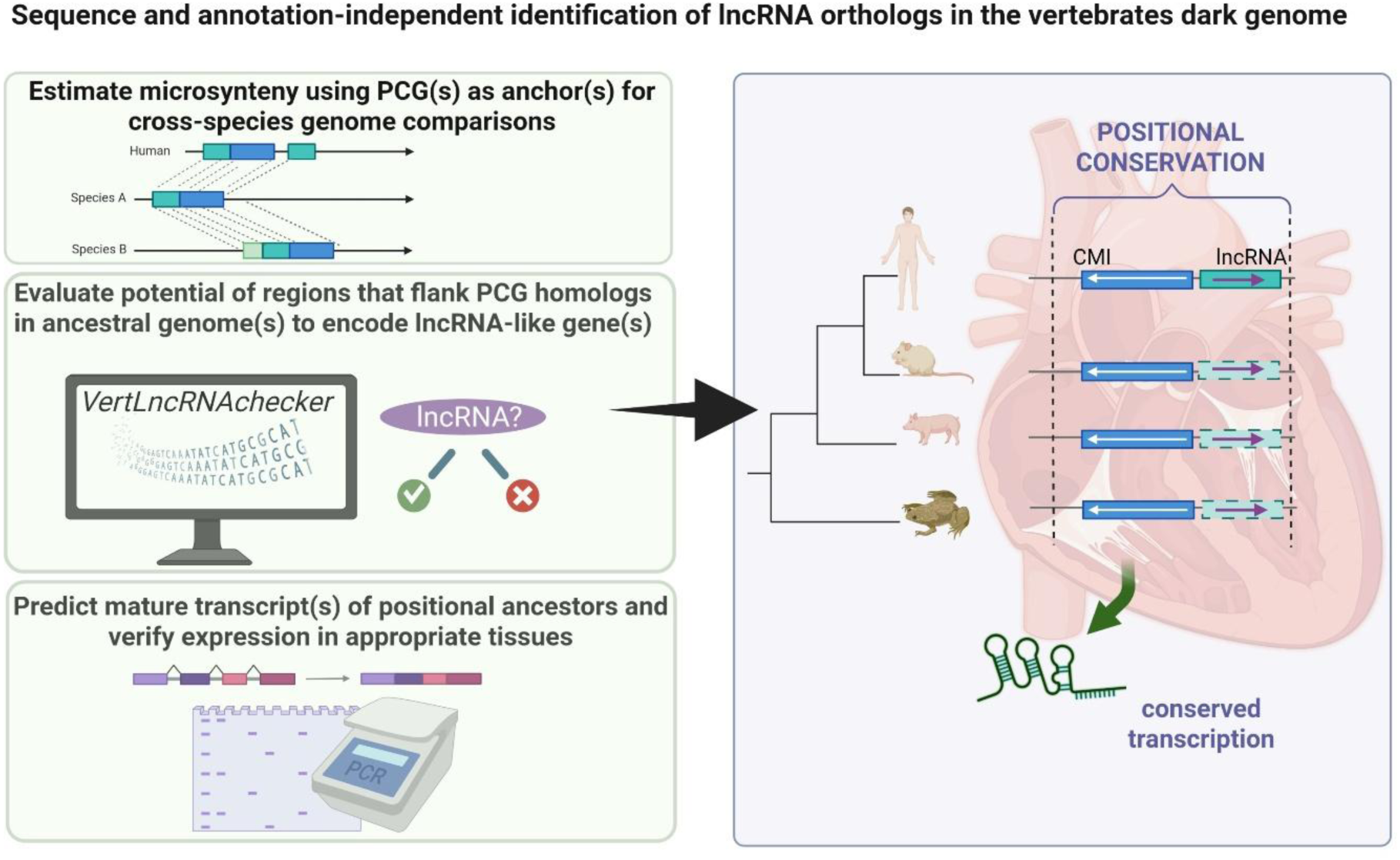

## BACKGROUND

Long non-coding RNA (lncRNA) genes are versatile regulators of genome function that produce transcripts ≥ 200 bps without encoding proteins [1]. They outnumber mRNAs, and they generally exhibit lower, spatiotemporally specific expression and markedly poorer sequence conservation than protein-coding genes (PCGs) [1]. They mediate diverse cellular processes by acting on local (*cis*) or distant (*trans*) genomic targets. They frequently regulate nearby PCGs, providing contextual and functional clues according to the ‘guilt-by-association’ principle [1, 2]. Their remarkable diversity of molecular mechanisms (comprehensively reviewed in [2]), inherently complicates their functional characterization, especially in complex vital organs, such as the heart. Cardiac lncRNAs regulate cell differentiation [3], function[4] and pathology [5, 6], and stress responses, such as tissue adaptation that compensates for myocardial injury [7]. Despite their elusive nature, they represent novel players in cardiovascular disease (CVD) and regenerative medicine[8, 9] that cannot be set aside in functional studies.

Because human cardiac tissue is scarce, the understanding of the role(s) of several genes in the heart relies heavily on animal models. The laboratory mouse is the gold standard [10], despite its evolutionary distance from humans which necessitates careful interpretation [11]. Pigs exhibit more similarities to humans than mice regarding heart physiology[12]. Evolutionarily distant vertebrates, like chickens[13], frogs,[14] and the zebrafish[15] occasionally unveil deeply conserved aspects. Nevertheless, the generally poor sequence homology of lncRNA genes introduces species specificities that limit in vivo experimentation, given that the key prerequisite for their usability is the existence of conserved orthologs of the human lncRNA genes of interest in these models. LncRNAs expressed in the human heart often appear to lack vertebrate counterparts, and vice versa. As a representative case, the murine *Hdnr* lncRNA gene, which is indispensable for mouse heart development [16], lacks human orthologs based on primary sequence homology thereby hindering clinically meaningful cross-species comparisons and translation to human pathology. More importantly, *Hdnr* highlights a paradox: how a gene essential for mouse cardiac function can be entirely absent in humans despite the highly conserved nature of mammalian heart development.

LncRNA conservation entails parameters besides primary sequence, such as structure and expression from syntenic loci[17]. LncRNA position relative to neighbouring genes emerges as a critical dimension of functional conservation. Amaral *et al.* first showed that the positions of the lncRNA gene promoters are preserved relative to the neighboring orthologous PCGs, to ensure that the lncRNA transcripts are co-expressed with their neighboring PCG products (mainly developmental transcription factors) in the same tissues[18]. Consistently, there are topologically conserved lncRNA[19] and miRNA genes[20, 21] that exert similar functions through interacting with the same genes in proximity, despite poor sequence homology. On a similar note, lncRNAs with conserved genomic locations and patterns of binding sites for RNA-binding proteins exist from zebrafish to human, which conserve their functions by interacting with similar proteins[22].

Positional conservation is prominent within vertebrates. Conserved blocks of genes occupying similar relative genomic positions (a phenomenon known as ‘synteny’) apply for hundreds of lncRNAs in amniotes and more distant vertebrates [18, 23, 24]. Such motifs are traced even earlier in evolution, as chordate lncRNAs frequently exhibit ‘microsynteny’, characterized by the preservation of small, localized genomic neighborhoods with conserved local gene order, transcriptional orientation, and relative spacing across different species, despite minimal or no lncRNA primary sequence conservation [25]. This pattern suggests that selective pressure acts primarily upon genomic context rather than nucleotide composition, implying functional links between lncRNAs and their adjacent coding or regulatory loci. Syntenic and microsyntenic blocks are likely inherited from a common ancestor and often encompass functionally related genes or regulatory elements involved in essential biological functions [26]. These observations highlight the maintenance of functional genomic associations during evolution and support the notion that genomic position is a key determinant of lncRNA function[18]. Consequently, a lack of sequence conservation does not necessarily negate the existence of lncRNA orthologs, as conserved sequence segments within syntenic regions can indicate the presence of lncRNA gene orthologs across species. To the best of our knowledge, no currently available incorporates positional conservation and microsynteny criteria to guide the identification of heart-related lncRNA gene orthologs, thereby limiting the effective use of vertebrate models in CVD research.

We hypothesized that the strong evolutionary constraints on the vertebrate hearts promote retention of genomic position and transcriptional orientation in heart-associated lncRNAs, particularly those acting as cis-regulators of indispensable neighboring PCG, irrespective of sequence similarity. This is supported by the finding that in mouse and chicken embryonic hearts, most cis-elements at the same genomic position remain functionally conserved at equivalent developmental stages, despite the lack of sequence conservation [27]. Encouraged by these findings, we developed an integrative cardio-informatics pipeline that combines phenotypic and single-cell sequencing data with synteny analysis and neural network-based modeling to predict evolutionarily related, including distant, heart-relevant lncRNA loci across vertebrate species commonly used in cardiovascular research.

Using this approach, we detected and experimentally verified a subset of lncRNA genes transcribed in mouse, pig, and frog hearts that occupy genomic positions corresponding to those of their human counterparts relative to neighboring PCGs. Of note, many of these previously unrecognized lncRNA-like loci reside within genomic regions currently unannotated or labelled as “junk DNA”. Furthermore, we developed a user-friendly web interface, *VertLncRNAchecker*, which discriminates lncRNA gene candidates from other non-coding genomic regions. Our workflow overcomes the limitations imposed by poor sequence conservation, evolutionary distance, and incomplete genome annotation. It not only represents a complementary resource to guide the functional lncRNA studies in vertebrate models of cardiovascular research, but due to its species-agnosticism and retrainability, it can be adapted to other research fields with similar dependence on animal models.

## MATERIALS AND METHODS

### Phenotype-driven identification of PCGs indispensable for cardiomyocyte development and function

To identify PCGs essential for cardiomyocyte differentiation and function (CM-Indispensable genes, CMIs), we screened the Mouse Genome Informatics database for candidates whose knockout leads to mouse phenotypes with relevant cardiac abnormalities. We downloaded the file ‘All Genotypes and Mammalian Phenotype Annotations (tab-delimited)’ and mined it for terms specific to cardiomyocyte differentiation, generating a list of CM-relevant mouse phenotype IDs. The list was used as an input in the field “Mouse phenotypes & mouse models of human disease” of “Genes and markers query”, selecting ‘protein coding gene’ as a ‘Feature type’. The resulting gene list was used as input in the ‘Batch query’ option, selecting ‘Mammalian Phenotype (MO)’, to match each gene with the corresponding phenotype IDs and descriptions. In this way, we retrieved 1828 CMIs, each assigned to their corresponding mouse cardiac phenotype(s). Their human orthologs were obtained from HGNC (https://www.genenames.org/). To infer the CMIs that are functionally conserved in humans, we subsequently used the 1828 genes as an input in the ‘Batch query’ option, selecting the output “Human Disease (DO)’ and identified all human diseases related to impairment of each gene. Then, we screened for those genes with cardiac involvement, that is, if impaired, they result in human cardiovascular diseases and/or in syndromes that include congenital heart defects. To systemically curate the syndromes with cardiac involvement, we used the GenMed database of NCBI. The 1828 genes, their corresponding mouse cardiac phenotypes, the associated human diseases, and their human homologues are listed in Supplementary Table S1.

### Single-cell RNA sequencing analysis

Single cell-RNA sequencing data from hearts from 18 human embryos (5-25 gestational weeks) [28] were downloaded from Gene Expression Omnibus (GSE106118) and processed by the R package Seurat (version 5.3.0). Quality control parameters, including the number of genes, transcripts per cell, and the fraction of transcripts from mitochondrial genes [29], were used to exclude low-quality cells (feature counts <100), potential doublets (feature counts >5,000), and damaged cells (>20% mitochondrial RNA). Transcripts expressed in ≤ 3 cells were removed to reduce low-abundance noise. For segregating early and late cell subpopulations and their features, the Seurat object included only the four developmentally earliest or latest subsets. The preprocessed dataset included 2627 cells and 22,582 genes. Gene expression was normalized using ‘LogNormalize’ and scaled with the ‘ScaleData’ function. Highly variable genes were identified using the ‘FindVariableFeatures’ function with the *vst* method. Principal component analysis (PCA) was performed by the ‘RunPCA’ function on the top 4000 most variable genes. For clustering, the top 15 principal components were implemented in Seurat via the ‘FindNeighbors’ and ‘FindClusters’ functions. Two-dimensional visualization was achieved using UMAP through the ‘RunUMAP’ function. For cluster annotation, we manually assigned clusters by combining three approaches: (a) characterization of cluster-specific gene expression patterns using the ‘FindAllMarkers’ function and application of the top 100 positive markers to identify associated biological functions and potential cell types by employing EnrichR; (b) identifying clusters enriched for individual markers using the ‘FeaturePlot’ and ‘VlnPlot’ functions; (c) including information about sample origin with respect to the developmental stage.

### Gene Set Enrichment Analysis, functional interaction networks, relative lncRNA-CMI mapping in the human genome

Gene Set Enrichment Analysis (GSEA) on CMIs, along with the protein-protein interaction networks of their products, were conducted as previously described [30, 31]. To analyze the topology and proximity of CMIs, date were extracted from the gff3 files Comprehensive gene annotation and Long non-coding RNA gene annotation available on genecodegenes.org (release versions 47) and the function “closestBed” of bedtools on the Galaxy server, based on our previous approach [1]. We filtered for CMI/lncRNA genes pairs with transcription start sites (TSSs) less than 5000 bp apart from each other. The PCG neighbors were integrated with the DEGs from scRNA-seq analysis to identify CMIs expressed during human heart development. The retrieved gene pairs were further classified according to the orientation of the transcripts (sense/antisense and intergenic, embedded, or overlapping). The naming schemes were standardized based on the most recent guidelines [32].

### Ortholog identification and prediction of microsynteny by database sequence similarity search

The nucleotide sequences corresponding to the exonic regions [33] of the human lncRNAs under investigation were retrieved from Ensembl (Release 112) [34], together with murine lncRNA sequences that are well-annotated but lack characterized human counterparts. The sequences were subsequently used as ‘probes’ in reciprocal BLASTn searches against the genomes of *Homo sapiens* (human), *Mus musculus* (Mouse), *Sus scrofa* (Pig), and *Xenopus tropicalis* (Frog) in Ensembl, with default parameters. The putative homologous sequences were evaluated based on the following criteria: a) best alignment hit defined by the lowest Expect (E) value (cut-off <0.01), b) sequence length greater than 200 bp, c) lack of protein-coding potential, assessed using the Coding Potential Calculator (CPC) [35] with default parameters, d) evidence of microsyntenic conservation [25, 33], by examining the chromosome localization and transcriptional orientation of the putative *lncRNA-like* sequences relative to their associated PCGs [18] using the Ensemble genome browser. For each mutually best match (i.e., vertebrate sequences exhibiting the highest degree of similarity to the query human lncRNA sequence than any other sequence), the flanking genomic regions stretching 5 kbps upstream and downstream of each hit were extracted. The intron-exon boundaries of the putative lncRNA sequences were predicted with SpliceBERT [36]. The PCG orthologs in the metazoan genomes under study were identified using their respective gene symbol in Ensembl [34]. For genes lacking annotation, the corresponding human PCG amino acid sequences were retrieved from the publicly accessible non-redundant sequence database NCBI’s RefSeq [37]; the longer and best-annotated protein isoform was chosen. These sequences were then used as ‘seeds’ to parse the metazoan genomes available in Ensembl [34], by reciprocal BLASTp [38] with default parameters. All sequences were retrieved, and all homology searches were performed independently by two authors (E.Y. and A.P.)

### Deep learning model for computational prediction of lncRNAs

A neural network model was developed based on diverse groups of human lncRNAs and applied across vertebrate species, in Python v3.13.0, following the steps below:

a. ***Acquisition of RNA sequences:*** Human ncRNA sequences were collected from the HUGO Gene Nomenclature Committee (HGNC; https://www.genenames.org/), last accession date January 10, 2025). The ‘positive dataset’ consisted of 3416 human lncRNA sequences. Other classes of ncRNAs, including miRNAs (microRNAs), tRNAs (transfer RNAs), rRNAs (ribosomal RNAs), snRNAs (small nuclear RNAs), snoRNAs (small nucleolar RNAs), vtRNAs (vault RNAs), RNY (Y RNA), and scaRNA (small Cajal body specific RNAs), comprised the ‘negative dataset’. PCG sequences were excluded from both positive and negative sets to prevent the model from exploiting coding-specific sequence features and to enforce learning from non-coding sequence characteristics, which differ substantially from those of PCGs. To minimize redundancy and sequence similarity bias, a non-redundant dataset containing sequences sharing less than 95% pairwise identity was compiled.
b. ***Feature Extraction:*** From each RNA sequence, 32 handcrafted features were extracted to capture both its compositional and structural characteristics. The sequences were transformed into numerical representations using one-hot encoding, where each nucleotide (A, T, C, G, U) was mapped to a 5-dimensional binary vector. To ensure uniform input dimensions for downstream neural network processing, the sequences were padded to a fixed maximum length using the *pad_sequences* function implemented in Keras.
c. ***Data Processing and Batching:*** The merged dataset was randomly partitioned into ‘training’ (80%) and ‘validation’ (20%) datasets, using stratified sampling to preserve class distributions. The pipelines were constructed using TensorFlow v2.16.1 *tf.data.Dataset* API, for batching, shuffling, and prefetching of data for GPU acceleration and scalable model training.
d. ***Model Architecture:*** The classification model was implemented using TensorFlow and the Keras Functional API. It relied on a dual-input (two-branch) neural network architecture. In the first branch, the one-hot encoded RNA sequences were processed through bidirectional LSTM (BiLSTM) layers to capture sequence dependencies and contextual patterns. In the second branch, the pre-extracted numerical features were fed into a series of fully connected (dense) layers. The outputs from both branches were concatenated and passed through additional dense layers; in the final output layer, a sigmoid activation function was applied to generate a binary classification score, indicating the likelihood that a given sequence corresponds to a lncRNA.
e. ***Model performance:*** To optimize performance and generalization: i) EarlyStopping was employed to prevent overfitting, ii) ModelCheckpoint was used to save the weights corresponding to the epoch with the highest validation accuracy, iii) due to class imbalance, class weighting was applied to penalize misclassification of minority samples and improve sensitivity for under-represented lncRNA sequences. The performance was evaluated on the validation dataset using the following standard classification metrics: Accuracy, Precision, Recall (Sensitivity), F1 Score, and AUC (Area Under the ROC Curve).
f. ***Cross-Species Model Application:*** To evaluate generalization across vertebrate species, the model - trained exclusively on human ncRNAs - was applied to sequences annotated as lncRNAs in Ensembl 112 from taxonomically diverse vertebrate species, including chimpanzee, macaque, mouse, pig, dog, chicken, zebrafish; frog was excluded due to the lack of annotated lncRNAs in Ensembl. Each sequence was transformed into 4-mers, and cosine similarity was calculated against the 4-mers of the known human lncRNA sequences. Vertebrate sequences exhibiting a maximum cosine similarity ≥80% were classified as true lncRNAs.
g. **Development of the *VertLncRNAchecker*:** A web application demonstrating the algorithm is available at https://vertlncrnachecker.web.app/. Users can upload either individual RNA or genomic sequence, or batch files (.csv or .xlsx) containing multiple sequences, to be classified as lncRNA or non-lncRNA. Optionally, users may provide a known homologous lncRNA sequence to optimize comparative analysis. All functionalities are available via a RESTful API at https://vertlncrnachecker.web.app/apidocs/ (access to the materials during the review process is available upon request)

### Acquisition of material from human heart tissues

Human heart tissues were obtained from 4 CVD patients (average age: 69.8 years) who underwent cardiac surgery at the University Medicine Rostock (Table 1). Tissue excision and subsequent use for research purposes were approved by the Ethics Committee of the Rostock University Medicine (A 2018-0037). Patients gave written informed consent pre-operatively, according to the Declaration of Helsinki. Excision was performed with a cut-and-sew method: the left atrial appendage was amputated at the base of the appendage by excision, and the opening was oversewn in multiple fashions (running/mattress sutures, single/double layer, with or without felt pledged reinforcement) by a non-absorbable suture. The tissue was immediately transferred into an ice-cold tube containing modified Tyrode’s solution (13.6mM NaCl, 5.4mM KCl, 0.9mM CaCl_2_, 1 mM MgCl_2_·6H_2_O, 0.33mM Na_2_HPO_4_, 30mM 2,3-butanedion monoxin, 5mM HEPES; pH 7.4) and stored at −80°C.

**Table 1:**
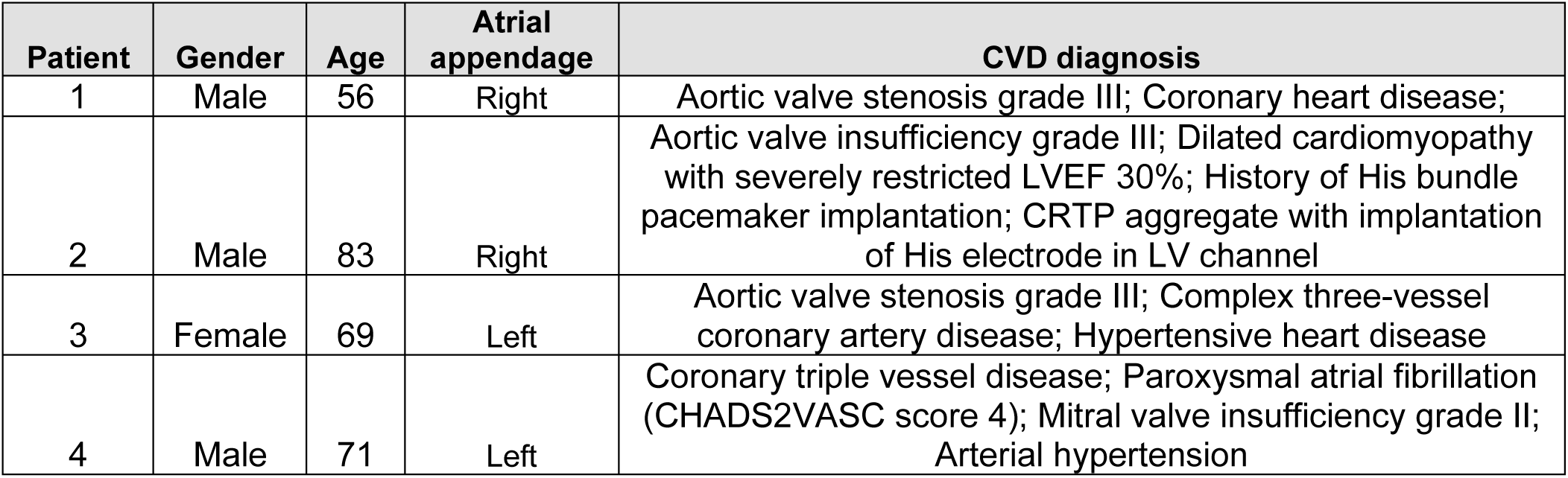
Overview of CVD patients from whom heart tissues were acquired.

| **Patient** | **Gender** | **Age** | **Atrial appendage** | **CVD diagnosis** |
| --- | --- | --- | --- | --- |
| 1 | Male | 56 | Right | Aortic valve stenosis grade III; Coronary heart disease; |
| 2 | Male | 83 | Right | Aortic valve insufficiency grade III; Dilated cardiomyopathy with severely restricted LVEF 30%; History of His bundle pacemaker implantation; CRTP aggregate with implantation of His electrode in LV channel |
| 3 | Female | 69 | Left | Aortic valve stenosis grade III; Complex three-vessel coronary artery disease; Hypertensive heart disease |
| 4 | Male | 71 | Left | Coronary triple vessel disease; Paroxysmal atrial fibrillation (CHADS2VASC score 4); Mitral valve insufficiency grade II; Arterial hypertension |

### Acquisition of material from vertebrate animal hearts

Whole hearts were harvested from adult female C57BL/6 mice (15 weeks old) following anesthesia induced by intraperitoneal injection of Pentobarbital (Narcoren; Boehringer Ingelheim, Germany; 50mg/kg body weight) and euthanasia by cervical dislocation. *Xenopus tropicalis* adults were sedated on ice for 45 minutes, decapitated, and dissected. Whole batrachian hearts were excised and shock-frozen in liquid nitrogen. Protocols of Xenopus care and experiments were approved by the Government of Oberbayern (ROB-55.2-2532.Vet_03-17-102). For porcine heart tissues, adults were anesthetized using electric pliers before euthanasia by exsanguination. Tissue samples of ∼0.5cm2 were excised from each heart chamber, pooled to avoid chamber-bias, and stored at −80°C. All animals used in this study were handled according to the guidelines from Directive 2010/63/EU of the European Parliament on the protection of animals used for scientific purposes and in compliance with the German Animal Welfare Act. Mice were housed in the animal facility of the Rostock University Medical Centre and maintained in specified pathogen-free conditions with food and water ad libitum, while pigs of the German Landrace were housed in the experimental animal facility at FBN Dummerstorf additionally complying with the Animal Protection Farm-Animal-Husbandry-Ordinance (German TierSchNutztV).

### RNA isolation, PCR detection, and amplicon sequencing

RNA was isolated from 50–100 mg of tissue using Trizol and subjected to reverse transcription using the GoScript™ Reverse Transcription System (Promega, Madison, Germany). For PCR primer design, the mature lncRNA sequences of all the microsyntenically conserved DNA regions were predicted with SpliceBERT [36] and the computationally predicted sequences were used as input in Primer3Plus [39]. Touchdown and, where applicable, nested PCR reactions were carried out in a Biometra TAdvanced Thermal Cycler (Analytik Jena, Jena, Germany) in duplicates. The detailed experimental conditions are provided in Supplementary File 1. Amplicon sizes and quality were inspected on a 2% agarose gel containing ethidium bromide (Roth, Karlsruhe, Germany). The products were purified using ExoSAP-IT Express PCR Product Cleanup (Applied Biosystems) and sequenced (QIAGEN Genomic Services, August 2025). Sequence identity between the obtained amplicons and the computationally predicted sequences was estimated using CLUSTALW [40], and coding potential was evaluated using CPC [35].

## RESULTS

### Design of a cardio-informatics workflow for sequence-independent identification of distant lncRNA orthologs across vertebrate hearts

Based on the ‘guilt-by-association’ principle, we assumed conservation of the cis-genomic position of lncRNAs relative to PCGs that are essential for normal cardiac phenotypes. The pipeline for the identification of lncRNA loci that may host putative orthologs of human heart-related lncRNA genes is illustrated in Figure 1. As described in the following sections, mouse cardiac phenotypes are integrated with scRNAseq from hearts of developing human embryos to identify PCGs crucial for myocardial development and function. The PCGS are paired with their neighbouring lncRNA genes in the human reference genome and syntenic analysis is performed. Regions in the vicinity of the vertebrate PCG orthologs are retrieved and evaluated for their potential to host a lncRNA gene using a retrainable neural network model. The computationally predicted mature RNA products of selected candidates guide the design of PCR primers for in situ detection of similar RNA transcripts in vertebrate heart samples, and the captured amplicons are verified by sequencing analyses.

**Figure 1:**
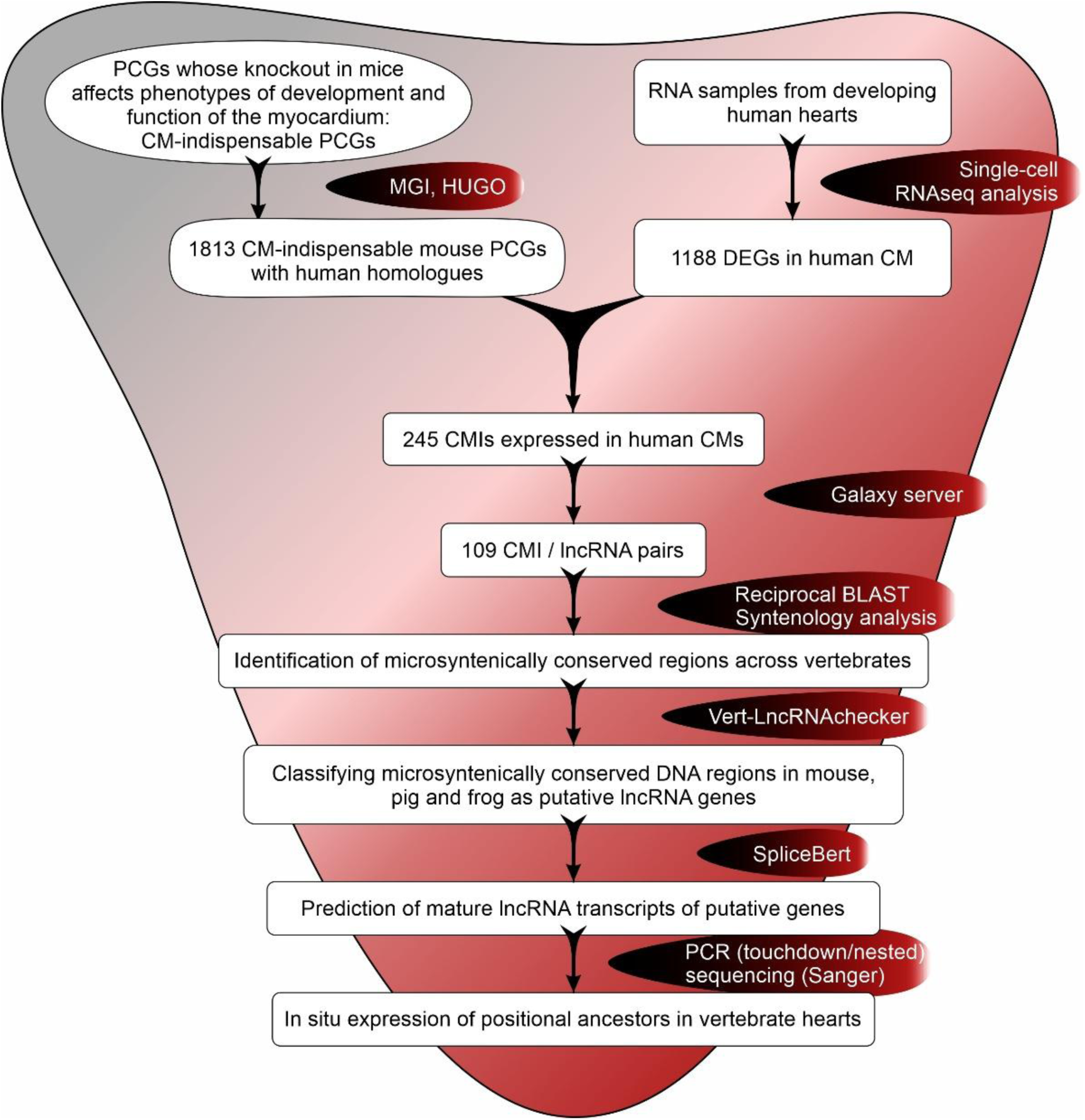
Cardio-informatics workflow. An overview of the steps designed and followed for the sequence-independent prediction and verification of distant orthologs of heart-related lncRNA genes. The workflow is versatile and can be adapted for (a) cardiac phenotypes of interest, by altering the input of knockout mouse phenotypes, (b) other differentiation stages, by altering the input of scRNAseq data, and (c) species of interest, by including the desirable species in the syntenology analysis and retraining the neural network model.

### CMI genes are expressed during human cardiomyocyte development and have functionally conserved human homologues

We systemically identified 1828 mouse PCGs whose knockout in mice establishes phenotypes of abnormal cardiomyocyte differentiation and function, with 1813 conserved human orthologs (Supplementary Table S1, referred to as CM-indispensable PCGs, CMIs). Of those, 11.6% (211/1813) have comparable heart-related functionality in humans and mice, meaning that their impairment in humans leads to heart diseases or syndromes encompassing congenital heart defects (Supplementary Table S1). Then, we inferred the CMIs implicated in myocardial development by identifying those that are differentially expressed in mature versus early human cardiomyocytes. To this end, we meta-analyzed the single-cell transcriptomics from cardiac cells of 18 human embryos, 5-25 gestational weeks old [28] (Figure 2A). Two clusters of ventricular cardiomyocytes (distinguished by cell distribution at early and late developmental phases were identified and compared (Figure 2B). The early cluster, representing immature ventricular CMs, is comprised of ∼96% cells from embryonal week 5-9, while the late cluster, representing mature ventricular CMs, is comprised of ∼78% cells from embryonal week 22-25. Overall, 1188 differentially expressed genes (DEGs) were identified (log2FC of −1.60 to 3.17, padj value ≤ 0.05). By overlapping them with the 1813 CMIs, we short-listed 245 CMIs undergoing transcript-level changes during human ventricular CM maturation (Supplementary Table S2). The cardiac roles of 58 of those are conserved in mouse and human (Figure 2C). The 245 CMIs are particularly enriched for cardiac morphogenesis and contraction (Figure 2D), contractile cardiac cell components (Figure 2E), and cardiac muscle contraction (Figure 2F), and their corresponding protein products form a highly interconnected gene regulatory network (Figure 2G). We inferred that these CMIs are the most essential for normal myocardial development and aimed to identify their human lncRNA gene neighbours.

**Figure 2:**
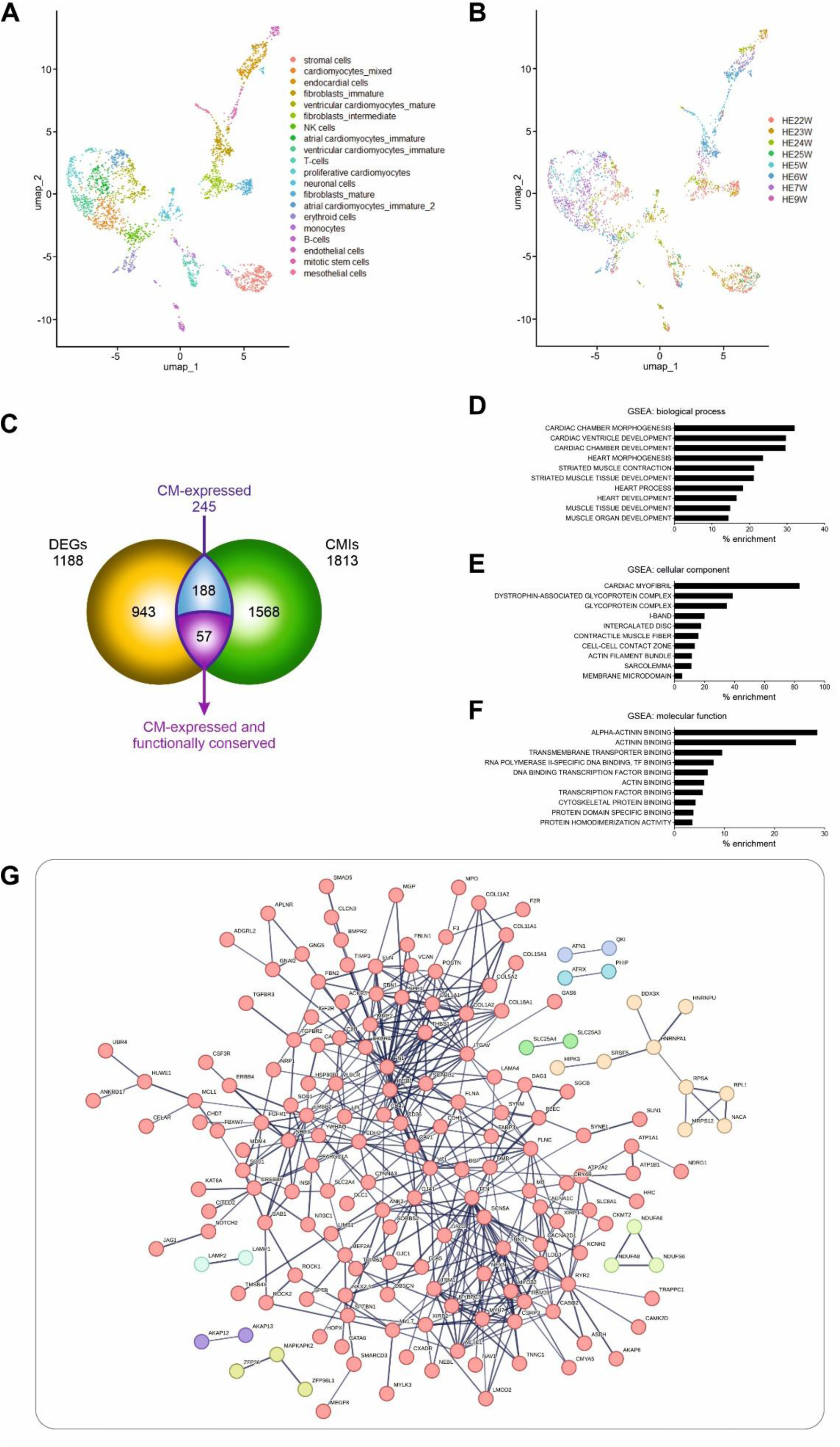
Computational identification and functional enrichment analysis of 245 CMIs expressed in the human heart. (A-B) UMAP representation of single-cell gene expression data for human embryonal (HE) cells by cell type (A) and developmental stage (embryonal weeks-W) (B). The two identified clusters of ventricular cardiomyocytes were classified to early, containing cells from embryonal week 5-7; and late, containing cells from embryonal week 22-25. (C) Pie chart depicting expression and functional conservation of CMIs. Of the 1813 human CMIs with vertebrate orthologs, 245 are expressed in human cardiomyocytes, while 57 are also functionally conserved in mouse and human. (D-F) GSEA for biological process (D), cellular component (E), and molecular function (F) of the 245 heart-expressed CMIs, (G) STRING network of the interactions among the protein products of the 245 genes.

### Human lncRNA genes adjacent to heart-expressed CMIs are microsyntenically conserved across vertebrates

We searched for annotated human lncRNA genes with a transcription start site (TSS) less than 5000 bp away from the identified 245 CMIs. Overall, 109 CMIs are flanked by lncRNA genes in the vicinity, at several gene arrangements and orientations. The most frequent orientation was the antisense (79%), and the most frequent arrangement was the overlapping (43%). There were also lncRNA genes in sense orientation (21%) and in embedded (28%), or intergenic arrangements (26%) (Figure 3A). Regarding the CMIs, 26 of them have conserved functions in human and mouse hearts. GSEA analysis for human disease phenotypes for this gene set indicated specific associations with lipodystrophy/lipoatrophy and cardiac abnormalities (Figure 3B). The remaining 83 CMIs are less enriched for diverse developmental disorders, implying more pleiotropic roles during human embryonic development (Figure 3C). We applied a series of prioritisation criteria to predict the most likely functionally related lncRNA neighbor(s) of the 109 CMIs. First, the nearest lncRNA within a 5000 bp window was selected, since spatial proximity is a critical factor in *cis*-regulation [41]. If a CMI had lncRNA genes in its vicinity, preference was given to the ones in the antisense direction, due to their well-documented roles in mediating cis-regulatory interactions [42, 43]. If more than one antisense lncRNAs was found within the 5 kb window, the annotated transcript was chosen. When no annotated antisense lncRNA was available, we considered nearby transcripts in opposite orientation relative to the CMI within the same genomic neighborhood, as well as the closest overlapping or directly flanking lncRNAs (e.g., divergent or sense-overlapping transcripts). This systematic curation of CMI/lncRNA pair genomic arrangements led to the identification of 110 unique human lncRNA genes associated with the 109 CMIs (Supplementary Table S3). To assess their conservation patterns across vertebrate genomes, we further selected the longest and best-annotated transcript for each locus from Ensembl. The entire sequences were used in reciprocal BLASTn searches against mouse, pig, and frog genomes, retaining only those hits that mapped to the same genomic position and with the same transcriptional orientation relative to the vertebrate CMIs. All candidate loci were subsequently inspected visually in Ensembl browser [37] for microsynteny across species (Supplementary Table S3).

**Figure 3:**
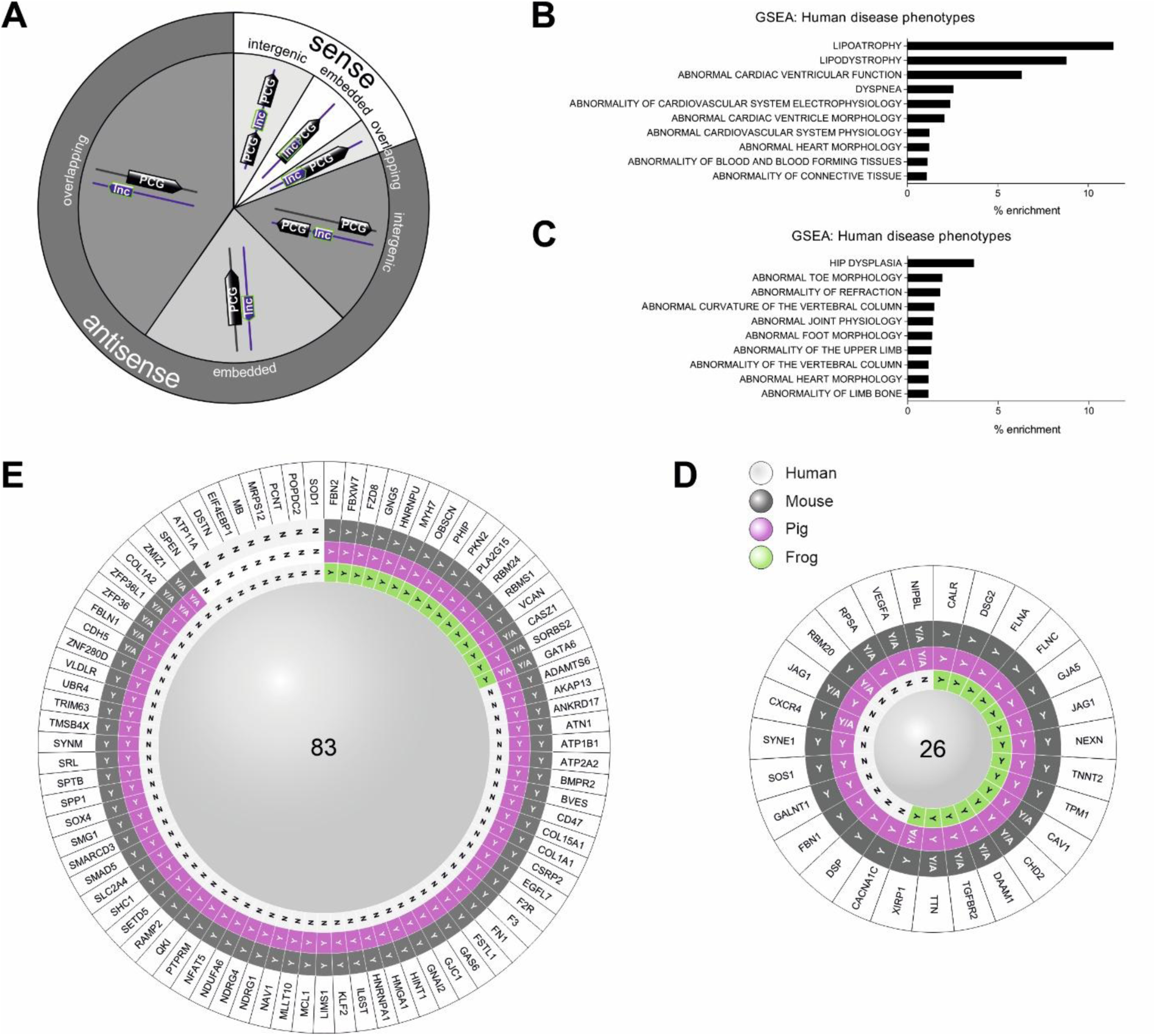
Genome mapping and microsynteny of CMIs and their neighbouring lncRNA genes across vertebrates. (A) Pie chart of the genome mapping of human lncRNA genes relative to CMIs ≥ 5 kbps upstream or downstream of the CMI transcription start sites (B-C) GSEA for human disease phenotypes of the heart-expressed CMIs with lncRNA neighbours for the 26 functionally conserved CMIs (B), and the remaining 83 CMIs (C). (D-E) Synteny analysis of the corresponding 26 (D) and 83 (E) CMI/lncRNA gene pairs. Annotations: A, lncRNA gene annotated in Ensemble; Y, syntenic conservation; Y/A, syntenic conservation in a region containing an annotated lncRNA gene; N, lack of syntenic conservation.

This approach revealed deep syntenic conservation from frogs to humans for one-third (31/110, 30.4%) of the analysed human lncRNAs. LncRNA genes near the 26 functionally conserved, heart-expressed CMIs showed particularly strong positional conservation: all of them were syntenically conserved in mammals (mice and pigs), while more than half (15/27, 55.56%) retained synteny in frogs (Figure 3D). Of the lncRNA genes near the remaining 83 heart-expressed CMIs, 74 (88.10%) and 73 (89.16%) showed synteny conservation in mouse and pig, respectively, whereas only 16 (19.28%) exhibited deep synteny in frogs; eight loci showed no detectable syntenic conservation across all tested species (Figure 3E). LncRNA microsynteny exhibited a continuous evolutionary pattern: once it emerged in a given lineage (amphibia, pig, or mouse), it was retained in all tested descendant species. Most of the syntenically conserved regions were not annotated, suggesting that still unidentified lncRNA orthologs may reside in the dark genomes of the vertebrates.

### *VertLncRNAchecker*: a neural network-based model for computational prediction of lncRNA genes in microsyntenically conserved regions

Syntenic conservation facilitates the identification of orthologous non-coding genes but is not absolute evidence for their existence[44]. To refine the accuracy of our workflow, we developed a neural network-based model that predicts whether a microsyntenic region of interest encodes a candidate lncRNA gene or not, through a binary classification process. We took into account that synteny-based comparisons depend on the presence of orthologous neighboring genes, perform optimally in evolutionarily closely related species, and become progressively ‘noisy’ as the phylogenetic distance increases [44]. Our model, which we termed *VertLncRNAchecker*, exploits datasets from vertebrate species that are commonly used in cardiovascular research and are evolutionarily related. The model incorporates a diverse set of features, including compositional, dinucleotide, and sequential properties, integrated using a hybrid BiLSTM architecture, and is available in our user-friendly web application (https://vertlncrnachecker.web.app).

First, we trained the model on the human genome, using multiple classes of well-annotated lncRNA sequences and evaluated its performance on an independent human validation dataset. Other classes of ncRNAs were used as negative controls. *VertLncRNAchec*ker distinguished human lncRNA genes from other ncRNA and/or genomic background (“junk”) DNA with high performance metrics: Accuracy (98.5%), Precision (99%), Recall (98.5%), F1 Score (98%), and AUC (0.9946). In a second step, the model was applied to annotated lncRNAs genes across reference vertebrate genomes relevant to cardiovascular research, namely mouse (*Mus musculus*), pig (*Sus scrofa*), chicken (*Gallus gallus*), and zebrafish (*Danio rerio*). To minimize evolutionary gaps among the selected species, we additionally included annotated lncRNA genes of intermediate relatives, namely primates (chimpanzee and macaque) and dog. Notably, the model robustly and accurately identified non-human lncRNA genes across all examined species (Table 2).

**Table 2.** Neural network model performance metrics (ranked by phylogenetic distance to humans).

|  |  | Vert-IncRNAchecker |  |  |  |  |  | Lncfinder | PreLnc |
| --- | --- | --- | --- | --- | --- | --- | --- | --- | --- |
| Organism | Annotated lncRNA genes(no.) | Accuracy | Precision | Recall | F1 score | AUC score | Detection percentage | Detection percentage | Detection percentage |
| Chimpanzee | 1778 | 0.984252 | 0.988656 | 0.995431 | 0.992032 | 0.893024 | 100% | 67.0% | 52.76% |
| Macaque | 4769 | 0.987209 | 0.989265 | 0.997877 | 0.993552 | 0.893201 | 100% | N/A | N/A |
| Mouse | 6305 | 0.979223 | 0.984026 | 0.994993 | 0.989479 | 0.866326 | 99.5% | 72.5% | 30.77% |
| Pig | 10971 | 0.994531 | 0.999908 | 0.994622 | 0.997258 | 0.994348 | 100% | N/A | N/A |
| Dog | 6479 | 0.972064 | 0.994629 | 0.977185 | 0.985829 | 0.85525 | 100% | N/A | N/A |
| Chicken | 11944 | 0.951105 | 0.963697 | 0.986103 | 0.974771 | 0.84989 | 100% | 67.0% | 54.01% |
| Zebrafish | 1492 | 0.891421 | 0.966887 | 0.918239 | 0.941935 | 0.765445 | 91.08% | 68.47% | 38.40% |

N/A: not applicable

Then, the model’s classification performance was compared against *LncFinder* [45] and *PreLnc* [46], two tools with well-demonstrated accuracy, robustness, and cross-species applicability. *VertLncRNAchecker* consistently recognized a higher percentage of lncRNA genes as ‘true’ in diverse vertebrate genomes, including chimpanzee, mouse, chicken, and zebrafish, relative to the comparators. Moreover, as *VertLlncRNAchecker* is organism-agnostic, it successfully classified annotated lncRNA genes as ‘true’ in species not included in the *LncFinder*, such as macaca, dog, and pig. For frog, performance evaluation could not be executed for any of the three tools due to the absence of annotated amphibian lncRNA genes in Ensembl. These metrics demonstrated the superior performance and suitability of *VertLncRNAchecker*. Subsequently, we applied *VertLncRNAchecker* to determine whether the the microsyntenically conserved regions (listed in Supplementary Table S3) harbor putative lncRNA genes. All examined regions were classified as *bona fide* lncRNA genes. We defined the microsyntenically conserved regions hosting computationally predicted lncRNA genes as positional ancestors of the human lncRNA genes and the respective CMI/lncRNAs as positionally conserved loci.

### Positional ancestors of human lncRNA genes are transcribed across vertebrate hearts

If the positional ancestors of human lncRNA genes are transcriptionally active, they should transcribe RNA molecules longer than 200 bps that lack protein-coding potential. Furthermore, if they are functionally analogous to their respective human lncRNA genes, they would at least be transcribed in the heart tissues of the corresponding species. To test these predictions, we selected four positionally conserved CMI/lncRNA loci for *in situ* experimental validations (Figure 4A). Two of them, namely FLNC/FLNC-AS1 and *TNNT2/ENSG00000286600*, showed deeply conserved syntenies from frogs to humans and entail heart-expressed CMIs with confirmed functional conservation in human and mouse. To avoid bias towards the subset of the 26 functionally conserved CMIs, we further included two positionally conserved loci from the larger subset of 83 CMIs from Fig. 3E, focusing on genes with well-characterized roles in the human heart. Thus, we selected GATA6/GATA6-AS1, where *GATA6* is essential for myocardial adaptation to pressure overload [47], while *GATA6-AS1* is implicated in aortic valve development [48]; and *ATP2A2/ENSG00000289311*, whereby *ATP2A2* regulates Ca^2+^ ventricular CM turnover during diastole [49]. The four selected CMIs regulate a broad range of heart molecular functions: the GATA6 protein is a transcriptional factor; the FLNC (filamin C) protein is a structural protein with striated muscle-specific expression, bridging the cytoskeleton with the extracellular matrix to maintain heart integrity and function [50]. The cardiac muscle troponin T, encoded by *TNNT2*, is a striated muscle component that binds tropomyosin and regulates muscle contraction [51]. The SERCA Ca^2+^-ATPase, synthesized by *ATP2A2*, is a membrane pump of the sarcoplasmic reticulum that restores cytosolic Ca^2+^ levels to their resting state for the next contraction [49]. Regarding their lncRNA counterparts, *FLNC* and *GATA6* overlap with the characterized antisense lncRNAs, while *ATP2A2* and *TNNT2* overlap with transcripts annotated in Ensembl as lncRNAs - in antisense and sense orientations, respectively – although the functions of these loci are currently uncharacterized in a cellular context (Figure 4A).

**Figure 4:**
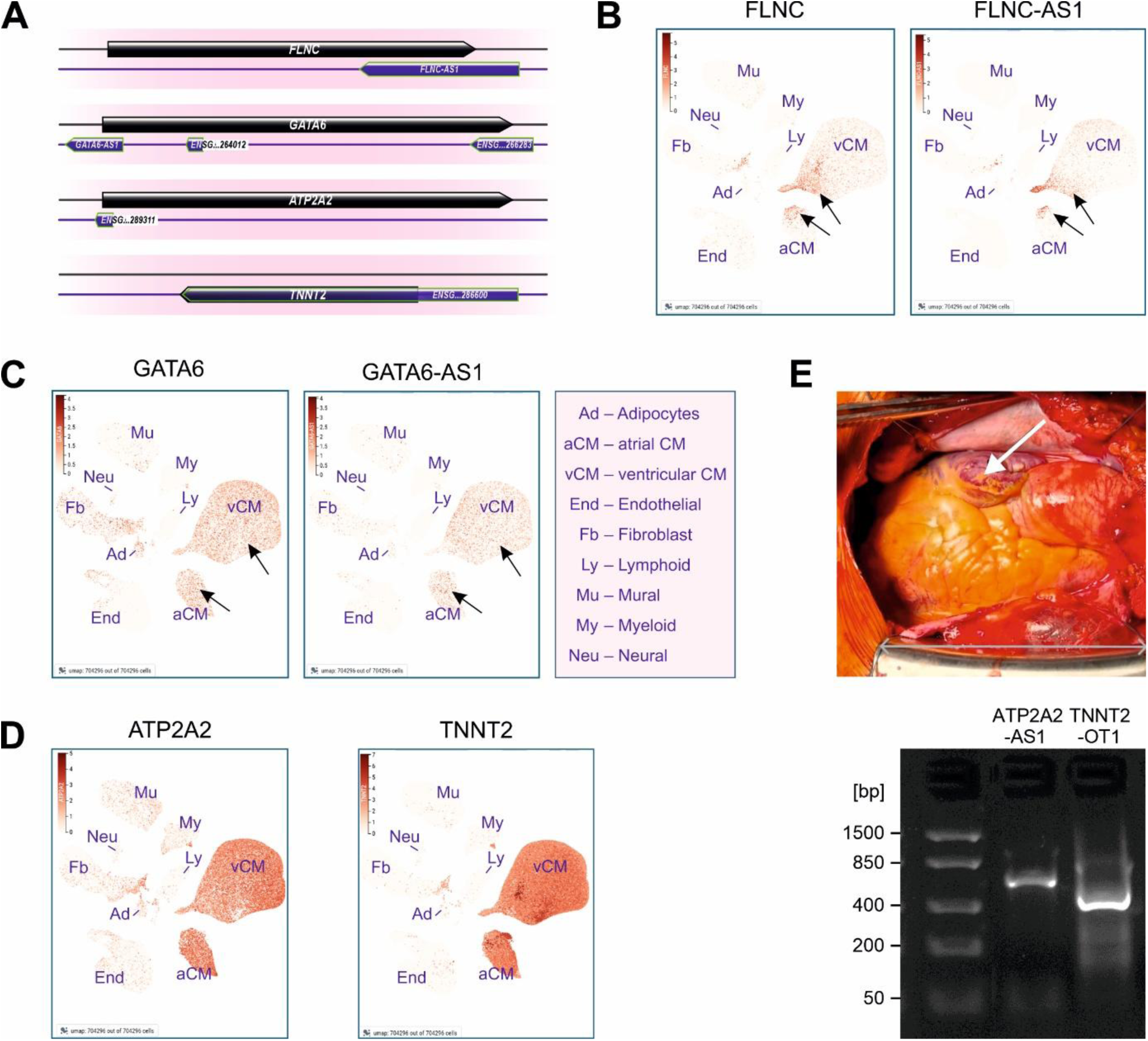
Expression of positionally conserved CMI/lncRNA pairs in the hearts of healthy human individuals and CVD patients. (A) Genomic arrangements of the four selected CMI/lncRNA gene pairs (B-C) Expression maps of the human FLNC/FLNC-AS1 (B) and GATA6/GATA6-AS1 (C) in the heart cell tumor atlas. The atlas contains transcriptomes from human hearts of 25 adults (14 females, 11 males, 20-75 years old) without a prior history of CVDs. The arrows indicate regions of co-expression of lncRNAs with the CMIs/lncRNA transcripts. (D) ATP2A2 (left) and TNNT2 (right) mRNA expression in the heart cell tumor atlas. (E) top: representative photo from the heart of a CVD patient undergoing surgery. The white arrow indicates the area from which the tested samples were obtained; bottom: PCR amplicons of the ATP2A2-AS1 and TNNT2-OT1 from RNA isolated from the retrieved patient heart tissues.

Expression was investigated in single-cell-nuclei expression data from healthy adult donors of the human heart cell atlas[52]; and in pathological samples from 4 CVD patients who underwent cardiac surgery (Table 1). FLNC-AS1 is normally expressed in cardiomyocytes across several regions, including atria, ventricles, interatrial septum, and apex, and co-expressed with FLNC transcripts, especially in ANV atrial cardiomyocytes (Figure 4B, see arrows). GATA6-AS1 is co-expressed with GATA6 (Figure 4C) in atria and ventricular cardiomyocytes. In patient samples, neither FLNC-AS1 nor GATA6-AS1 transcripts were detected. ATP2A2 and TNNT2 transcripts are highly expressed in atrial and ventricular cardiomyocytes of normal hearts (Figure 4D). Although the expression of their uncharacterized neighboring lncRNAs (ENSG00000286600 and ENSG00000289311) is non-minable from the heart atlas, we were able to detect their RNA transcripts (hereafter termed as ATP2-AS1 and TNNT2-OT1, respectively) in all pathological cardiac samples. The amplicons of ATP2-AS1 and TNNT2-OT1 were, correspondingly, 100% and 89% identical to the predicted mature lncRNAs (Figure 4E). Overall, the lncRNA neighbors of all selected CMIs are expressed in normal or diseased human hearts (Table 3).

**Table 3:** Expression of transcripts of selected lncRNAs in hearts of healthy individuals and CVD patients.

| **CMI** | **Human disease related with the CMI (MGI database)** | **lncRNA neighbor** | **lncRNA expression in normal hearts** | **No. of CVD patients detected** | **Identity<sup>b</sup>** | **Difference between the detected relative to the predicted mature transcript** |
| --- | --- | --- | --- | --- | --- | --- |
| FLNC | myofibrillar myopathy 5 | FLNC-AS1 | Yes | 0/4 | — |  |
| TNNT2 | dilated cardiomyopathy 1D; dilated cardiomyopathy 1D; hypertrophic cardiomyopathy 2 | TNNT2-OT1 | N/A | 4/4 | 100% |  |
| ATP2A2 | — | ATP2A2-AS1 | N/A | 4/4 | 89% | 64-bp truncation |
| GATA6 | — | GATA6-AS1 | Yes | 0/4 | — |  |

<sup>a</sup> Ratio of successful PCR amplifications in n out of four samples analyzed
<sup>b</sup> Nucleotide comparison between the predicted lncRNA fragment and the consensus sequence obtained by Sanger sequencing; coverage differences were considered. Identity of detected to predicted lncRNA transcript
N/A: not applicable

Subsequently, we investigated expression of the putative positional ancestors of the selected human lncRNA genes in vertebrate hearts. We predicted the corresponding mature transcripts in mouse, pig, and frog (Supplementary File 2), designed species-specific primers, and subjected heart tissues of the selected species to PCR analyses. The results are summarized in Table 4. All candidate transcripts were detected in pigs, with a high identity between the *in silico*-predicted and the *in situ*-detected sequences, ranging from 100% to a minimum 59% (Figure 5A). Even in cases of partial identity, the amplified sequences retained 100% identity within the corresponding regions of the predicted transcripts (Figure 5B-C). In mice, we verified expression of positional ancestors for *GATA6-AS1*, *FLNC-AS1, and TNNT2-OT1,* which are, correspondingly, 100%, 100%, and 82% identical to their predicted sequences. However, *ATP2A2-AS1* was not amplified in the tested mouse samples. In frogs, we captured a 120-bp amplicon of FLNC-AS1 as a band of approximately 800–900 bp (Table 4 and Figure 5D-E). Analyses of the *in situ*-detected RNA sequences using CPC v2.0 showed that all these products lack coding potential and therefore may represent lncRNA molecules. The respective three-dimensional (3D) structures of the PCR amplicons, generated using 3dRNA (http://biophy.hust.edu.cn/new/3dRNA), displayed typical lncRNA 3D motifs, such as stem-loops, internal loops and bulges, and multibranch (junction) loops, hierarchically organized into modular structural domains, a hint for lncRNA functionality[2, 53] (Supplementary Figure 1). Collectively, positional ancestors of all tested human lncRNA genes are expressed in mammalian hearts. For one of them, FLNC-AS1, a positional ancestor actively synthesizing lncRNA-like transcripts was detected in the three-chambered batrachian hearts, providing signs of conservation beyond the four-chambered mammalian heart.

**Figure 5:**
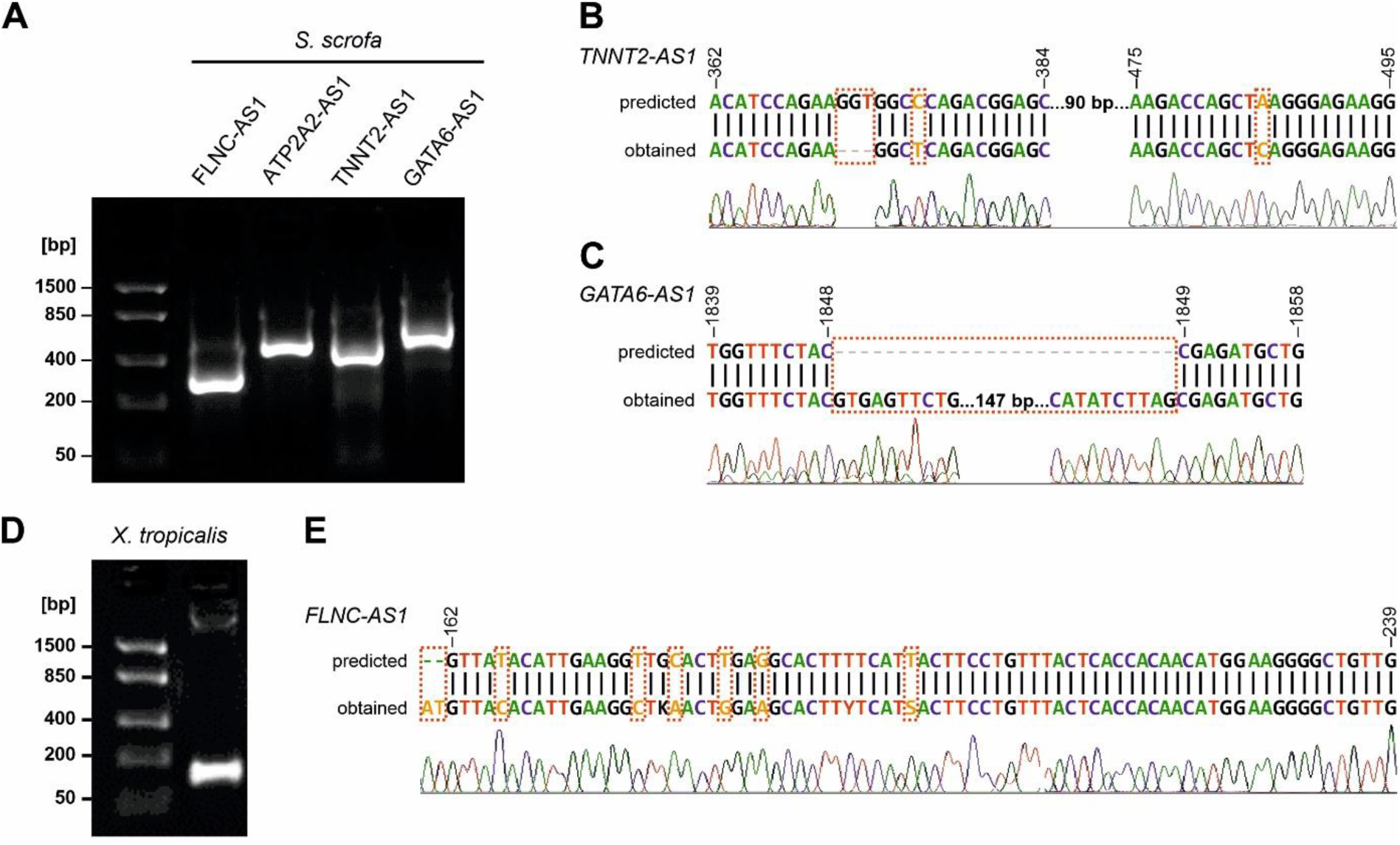
Positionally conserved lncRNAs are transcribed in vertebrate hearts. (A) Agarose-gel electrophoresis visualising PCR products for *FLNC-AS1*, *ATP2A2-AS1*, *TNNT2-OT1,* and *GATA6-AS1* in Sus scrofa hearts. Lane 1: DNA ladder. The predicted (upper row) sequences of the porcine (B) *TNNT2-OT1* and (C) *GATA6-AS1* lncRNA were compared to the corresponding experimentally obtained sequences (lower row). Orange characters indicate nucleotide substitutions, while grey characters indicate deletions in the predicted sequence versus the sequenced amplicon. Numbers indicate the nucleotide positions on the predicted lncRNA sequence. (D) PCR for *FLNC-AS1 in Xenopus tropicalis* heart samples (E) alignment between predicted and obtained sequences.

**Table 4:**
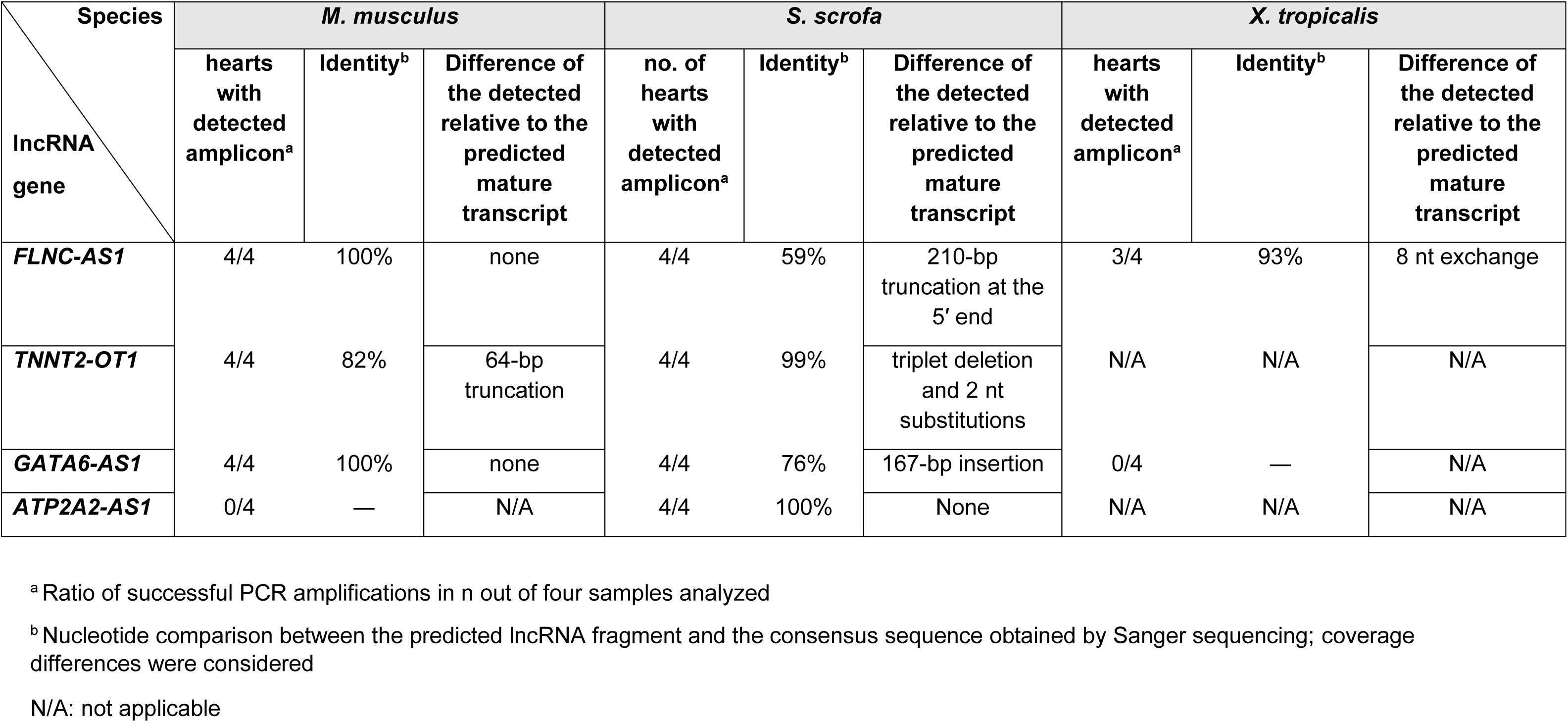
Expression of transcripts of positional ancestors in vertebrates relevant for cardiovascular research.

| Species<br>IncRNA<br>gene | *M. musculus* |  |  | *S. scrofa* |  |  | *X. tropicalis* |  |  |
| --- | --- | --- | --- | --- | --- | --- | --- | --- | --- |
|  | hearts<br>with<br>detected<br>amplicon<sup>a</sup> | Identity<sup>b</sup> | Difference of<br>the detected<br>relative to the<br>predicted<br>mature<br>transcript | no. of<br>hearts<br>with<br>detected<br>amplicon<sup>a</sup> | Identity<sup>b</sup> | Difference of<br>the detected<br>relative to the<br>predicted<br>mature<br>transcript | hearts<br>with<br>detected<br>amplicon<sup>a</sup> | Identity<sup>b</sup> | Difference of<br>the detected<br>relative to the<br>predicted<br>mature<br>transcript |
| ***FLNC-AS1*** | 4/4 | 100% | none | 4/4 | 59% | 210-bp<br>truncation at the<br>5' end | 3/4 | 93% | 8 nt exchange |
| ***TNNT2-OT1*** | 4/4 | 82% | 64-bp<br>truncation | 4/4 | 99% | triplet deletion<br>and 2 nt<br>substitutions | N/A | N/A | N/A |
| ***GATA6-AS1*** | 4/4 | 100% | none | 4/4 | 76% | 167-bp insertion | 0/4 | — | N/A |
| ***ATP2A2-AS1*** | 0/4 | — | N/A | 4/4 | 100% | None | N/A | N/A | N/A |

<sup>a</sup> Ratio of successful PCR amplifications in n out of four samples analyzed
<sup>b</sup> Nucleotide comparison between the predicted IncRNA fragment and the consensus sequence obtained by Sanger sequencing; coverage differences were considered
N/A: not applicable

## DISCUSSION

We present a cardio-informatics workflow detecting putative vertebrate orthologs of human lncRNA genes, independent of sequence conservation and genome annotation. It integrates heart developmental phenotypes, single-cell RNAseq data from developing hearts, synteny features, and a neural network model that predicts whether a non-annotated genomic region entails a putative lncRNA. We used CMIs as anchors for cross-species phylogenetic comparisons and termed as “positional ancestors” their flanking genomic regions across vertebrates that exhibit the same transcriptional orientation and lncRNA-like features as their human counterparts. For selected human lncRNAs, we validated that positional ancestors synthesize mature non-coding RNAs ≥200bps in the hearts of corresponding mammalian species. One positional ancestor was transcriptionally active outside the mammalian clade. Hence, the relative genomic position and transcriptional activity of the human lncRNA genes are retained in vertebrate genomes, even when corresponding sequences are highly divergent. Moreover, this pipeline facilitated the detection of potential lncRNA genes within the dark genome of vertebrate species. Whether the identified vertebrate transcripts exhibit similar biological functions to the prototype human lncRNAs remains an open question. Considering that the heart is a strict functional setting, where aimless gene changes could impact survivability and fitness, we envisage that the conserved transcription of positional ancestors across the hearts of all tested species is not random. Follow-up mechanistic investigation can shed light on their roles in mammalian heart homeostasis.

The prediction of lncRNA loci across diverse species genomes remains challenging. Robust computational tools (e.g., CPC [35] and CPAT [54]) combine sequence-derived features and machine learning to assess the coding probability of potential lncRNAs. However, reliance on coding potential alone may result in inaccurate classification, because a proportion of lncRNAs display features similar to protein-coding sequences [55]. Inadvertent biases are expected, considering that 5% of conserved mammalian lncRNAs are derived from lost PCGs [56]. Dependence of tools on genome sequence and annotation increases the risk of false-negative predictions, especially in species with incompletely annotated genomes. Although model re-training increases performance, only a few models are retrainable, and even so, their performance may be unstable for species whose genomes are partially sequenced or have variable compositions across organisms [45]. Tools exploiting features besides coding probability perform better but lack certain scalability and versatility [46]. For instance, *PreLnc* uses high-confidence lncRNA and mRNA transcripts to build prediction models through the selection of 11 features, including basic sequence, open reading frame, coding sequence, peptide characteristics, and functional definitions [46]. It performs well on *Homo sapiens* and *Arabidopsis thaliana* but has lower predictive power for mice [46]. *Lncfinder*, which exploits sequence intrinsic composition, structural information, and physicochemical properties, is more accurate for mouse datasets [45].

To overcome these limitations, we developed *VertLncRNAchecker*, a deep learning-based framework that integrates compositional, dinucleotide, and sequential features using hybrid bidirectional LSTM architecture. It accepts FASTA genomic or RNA sequence sequences from diverse data sources and rapidly returns binary predictions regarding their potential classification as lncRNAs. The tool demonstrated favorable classification performance across diverse vertebrate datasets, as well as higher predictive power *versus Lncfinder* and *PreLnc*. It is tailored for vertebrate species that are evolutionarily closer and share functional similarities in heart development and function, minimizing potential noise associated with broader phylogenetic distances [44]. The architecture of *VertLncRNAchecker* is retrainable, allowing future adaptation to additional species with shared ancestry and/or functional traits of interest. *VertLncRNAchecker* is available both as a standalone code and as a user-friendly website, maximizing accessibility by both computationally-savvy and -unsavvy users. It accepts both genomic and RNA sequence inputs to rapidly detect putative lncRNA sequences in diverse data sources.

The *in silico* predicted lncRNA transcripts from syntenically conserved regions guided primer design for capturing the products of positional ancestors across several vertebrate heart tissues. The fact that some detected amplicons were not 100% identical with their computationally predicted counterparts could reflect the presence of alternative-spliced variants. For predicting the exon splicing sites of the lncRNA genes, we used SpliceBERT [36] which is customized for pre-mRNAs. Nevertheless, pre-lncRNAs are spliced less inefficiently than pre-mRNAs, generating more transcript variants [57]. It is suggested that a certain level of elongation followed by early transcription termination reflects high regulatory complexity rather than splicing errors. Mechanistically, inefficient lncRNA splicing is linked to WDR82, a mammalian regulatory factor that, together with the ZC3H4 protein, promotes early transcription termination. Transcripts sensitive to WDR82 depletion are spliced less efficiently, and their mature forms extend beyond the 3′ end [57]. Accordingly, these small insertions and deletions within the identified discrepant amplicons may be attributed to weak splicing signals that increase transcript heterogeneity. There is a need for need for specialized exon-intron boundary prediction models, optimized specifically for lncRNA sequences. Such tools could integrate more effectively lncRNA-specific biological features to improve the detection of splicing sites and exonic boundaries.

Regarding the species examined, we found that the prediction efficiency was maximum for the pig genome, whereby all four positional ancestors were detected in every tested heart sample. The recent annotation of the pig cardiac transcriptome [58] further facilitates the use of our pipeline to discover the heart-relative lncRNAome. Given that porcine cardiovascular models have already been used to investigate lncRNA function *in vivo [59, 60]*, they represent a suitable system for mechanistic and functional studies and, eventually, translational relevance to human heart physiology and pathology. Since lncRNAs are inherently lowly expressed, higher amounts of heart tissue are required for their identification, which can be acquired due to the large size of porcine hearts. In mice, three out of the four selected human lncRNA genes exhibited positionally conserved orthologs, while the predicted transcript for the murine *ATP2A2-AS1* was not detected. There are several potential reasons for this lack of detection of the mature transcript in the tested material. First, the heart samples were retrieved from adult female mice, while the expression of this lncRNA positional ancestor may be either restricted to a specific cell type or developmental stage [61] that is not represented in the available tissues, or subjected to sex-specific regulation [62]. Second, the *ATP2A2-AS1* transcript could be an unnecessary by-product of the *cis*-regulation, following a well-recognized RNA-independent mechanism, whereby the lncRNA gene acts via its transcription itself, whereas the actual lncRNA product is unstable [63]. Furthermore, *GATA6-AS1* and *FLNC-AS1* expression was detectable in the healthy heart cell atlas but undetectable in human patient samples. It would be interesting to investigate whether their absence in the patient-derived tissue samples is causally associated with relevant cardiovascular pathologies.

We propose a theoretical framework to explain the existence of positional ancestors, by extrapolating the ‘guilt-by-association’ principle to an evolutionary context. The lncRNA homologues interacting with the same proteins may be under immense evolutionary pressure to preserve these associations within a syntenic locus, even when the primary sequence evolves at different rates. Resilience of the interaction of a certain lncRNA with its neighboring PCG, despite evolutionary changes in the lncRNA nucleotide sequence, is feasible through secondary or tertiary RNA structures [64]. Three-dimensional structures of lncRNAs are likely preserved throughout evolution, along with functional “modules”, interspersed with longer and less conserved stretches of nucleotide sequences [65]. Hence, if an alteration in the sequence of the positional ancestor is major but the resulting RNA product is re-composited to form structures still able to mediate the same RNA-protein interactions, then the functionality of CMI/lncRNA loci is retained despite the significant nucleotide changes. Looping within an RNA molecule creates binding modules that are not evident by primary sequence but could still mediate the same RNA-protein interactions [66]. These binding modules might suffice for preserving functional associations with certain proteins, given that RNA-binding proteins are selective for RNA hairpins or structures, not for primary sequences [67]. SLC16A1-AS1, a lncRNA which binds to E2F1 to co-activate its neighboring *SLC16A1*, represents an example that depends more on its loop-forming ability than its primary sequence for maintaining its cis-regulatory function [1]. Surprisingly, *SLC16A1-AS1* deletion mutants for several sequences did not lose their ability to interact with E2F1. Instead, some of them acquired thermodynamically more stable structures than the wild-type lncRNA, leading to stronger binding and efficacy of the co-activator complex [1]. Conversely, mutations in the promoter of a lncRNA gene could disrupt its co-transcription with important PCG neighbors, and the asynchronous transcription may render the PCG-lncRNA interaction no longer possible within a particular cell context, violating the ‘guilt by association’ principle This mechanism would explain the long-standing axiom that lncRNA genes have more conserved promoters than exon sequences [68]. It is also relevant in cases of transcript-independent mode of action of lncRNA genes, whereby the transcription of the lncRNA would be under higher pressure to be conserved than the actual RNA product.

Genetically engineered animal models link lncRNA genotypes to phenotypes. However, *in vivo* lncRNA knockouts can frequently lead to the absence of observable phenotypes [69], eventually rendering such studies cost-inefficient and impractical. Our workflow mitigates this experimental design risk by prioritizing lncRNAs with positional ancestors, the perturbation of which is more likely to yield clear phenotypic outcomes. whose perturbation is more likely to yield clear cardiac phenotypic outcomes. Future studies investigating whether positional ancestors flanking CMI orthologs retain their *cis*-regulatory roles, even when sequence conservation is low, could provide significant insights into lncRNA biology. Such studies are particularly timely, as they may shed light on lncRNA-mediated mechanisms of heart pathology and, at the dawn of RNA-centric drugs [70], inform the development of novel CVD therapeutics.

## CONCLUSIONS

We provide a systematic pipeline to prioritize non-coding genomic loci and their associated non-coding variants as candidate functional ancestors of human lncRNAs. By applying it to the evolutionarily constrained setting of the heart, we showed that transcriptionally active non-coding loci across vertebrate genomes occupy positions analogous to those of human lncRNAs relative to PCGs that are essential for cardiac function. Such lncRNA-like genes maintain their relative genomic position and transcriptional activity across vertebrate hearts, even when the corresponding sequences are highly divergent from the human lncRNA gene. The transcripts of the positionally conserved ancestors possess characteristic secondary structural modules, suggesting that disease-associated non-coding variants within these regions may disrupt conserved RNA structure and/or function. Therefore, these loci pose as strong candidates for conserved functional counterparts of the human lncRNAs. Given that the heart is a highly controlled functional entity, it is rather unlikely that the conserved transcription of positional ancestors across the hearts of all tested species is random and merits further investigation regarding its role in vertebrate heart homeostasis. This work fills a major evolutionary gap and provides a solid foundation for follow-up comparative mechanistic studies, cross-species functional validation, and the functional annotation of non-coding variants that are linked to cardiovascular traits. Genetic knockouts of positional ancestors in the hearts of not only the gold standard mice but also promising models like pigs and frogs can determine functional homology. Our approach informs targeted cross-species experimentation to determine whether these ancestral forms retain their regulatory interactions on the neighboring CMIs, contributing to conservation of myocardial phenotypes.

## DECLARATIONS

### Ethics approval and consent to participate

Human heart tissues were obtained from patients of the University Medicine Rostock Heart tissue excision, and subsequent use for research purposes was approved by the Ethics Committee of the Rostock University Medicine (A 2018-0037). Patients gave written informed consent pre-operatively, according to the Declaration of Helsinki. Mice were kept in the associated Central Laboratory Animal Facility in accordance with the German Animal Welfare Act. Pigs of the German Landrace were housed in the experimental animal facility at FBN Dummerstorf in compliance with European regulations for husbandry and experimental conditions and the national Animal Welfare Act and Animal Protection Farm-Animal-Husbandry-Ordinance (German TierSchNutztV). *Xenopus tropicalis* hearts were provided by the Biocampus Martinsried of the Ludwig-Maximilians-University, Munich.

### Consent for publication

All authors have read and consent to the final version of the manuscript.

### Availability of data and materials

The publicly available scRNA-seq data analyzed in this study were download from GEO (accession number GSE106118). All other data and methods will be made available upon request for those aiming to reproduce results or replicate procedures.

### Funding

RD acknowledges funding by the Deutsche Forschungsgemeinschaft (DFG) for projects GRK 2901/1– 501988175 and DA 1296/10-1 – 572594361; the German Federal Ministry for Economic Affairs and Climate Action (16KN083635); and the German Federal Office for Agriculture and Food (281A819B21). SL acknowledges DFG, project LO 3128/4-1 – 561811975.

### Authorś contributions

**AMG:** Conceptualization, Data curation, Formal analysis, Investigation, Methodology, Writing – original draft, Writing – review and editing; **AP:** Conceptualization, Data curation, Formal analysis, Investigation, Methodology, Writing – original draft, Writing – review and editing; **EY**: Software, Investigation; **SK:** Resources, Software**; SM:** Methodology, Data curation, Visualization, **JS:** Resources; **AR:** Formal analysis, Investigation, Methodology, Writing – original draft, Writing – review and editing, **OC:** Investigation; **CN:** Investigation; **CE:** Resources; **RD:** Conceptualization, Funding Acquisition, Resources, Writing – review and editing, Supervision; **SL:** Conceptualization, Data curation, Formal analysis, Investigation, Methodology, Visualization, Writing – original draft, Writing – review and editing, Supervision

## Acknowledgements

We are grateful to Madeleine Bartsch for the isolation of frog hearts; Luong Chau for supporting RNA isolation; and Franziska Witt for technical support with PCR.

## Notes

### Competing Interest Statement

The authors have declared no competing interest.

