## Supplementary material for "An integrative cardio-informatics approach identifies positionally conserved ancestors of human lncRNA genes transcribed across vertebrate hearts": Supplementary File 1.docx

**Predicted transcripts of selected heart-related lncRNA genes and their positional ancestors in mouse, pig and frog.**

The spliced RNA products of selected positional ancestors were predicted using SpliceBert. The yellow highlighted sequences are the ones with the highest similarity as compared (and therefore suspected to harbor conserved domains/interfaces for interactions with proteins)

1. **LOCUS OF INTEREST: FLNC/FLNC-AS1**

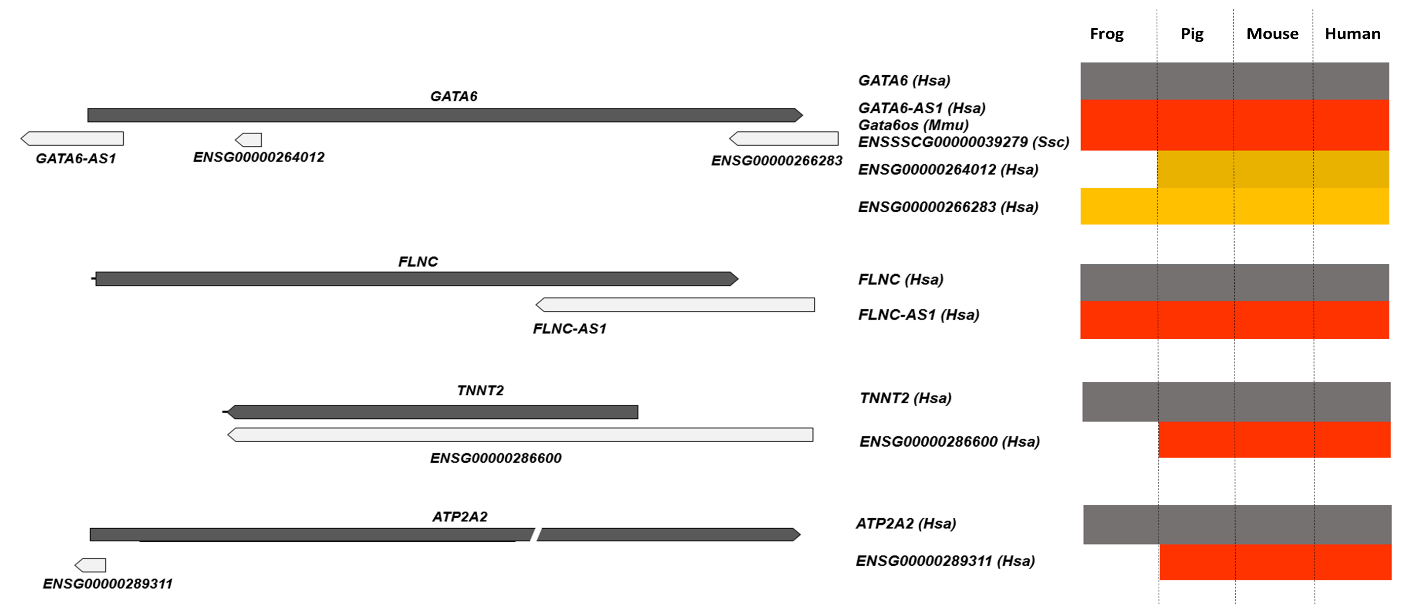
 frog pig	mouse human

**PREDICTED PRODUCTS**

**>Human_FLNC-AS1**

ATTCGTTCAACAAGCGCTTTGGACCTGACGCTGTTAGACGCCGAGACCCCAGAGAAGAGA

CGCAGTTACTCATCTCAAGGGCAGGGTCTGGGTCTTTATCGTACTGCAGGCTCCACCCAG

AATGGGCACCGATGTGTGTAGTGACCACTAACAAAAATGGCTCAGTGGGAGGAGCGGGGG

CACCACGGGACAGGCGGTCTGGAAAATTCCCTAAGGTGATCTCTGCCTTGGATGGGCCCT

CCACGGCCAGTGACAGACCCCCTGTAGGTGGAGGTGTGGCCAAGATCAGAGGTCCTGCCC

TCACCCCTGCCCCTGGCACCACTGGCTCCTCTGTGGCCTGCAGCAGGTGAGAGGGAACAG

TCAGTGACTGGGGAAGGCCCAGGAGTTTGACCCATGCTATGACTGAAGCTGGAGAGTGAG

GAAAGAACAAGGTAGTTAATCATGAAACCAGCGTGGCAGGAGAGGTGCGTCAGCTGCAAG

AGAAGGAGTGCTGTGCCCAAGTGGGGTTCCATCTGCCCAGCCTAACTCTTGCCTGCTCCC

CAAGAGGCCCCCAGACCAT

>**Mouse_ predicted**

CCTGGAGCCAGAGTACTCTAGAATGAGCCCTTGGGATCCCTGCAGCCTCCCAACGTGCAG

GGAGGCAGCGCAGCTCCGCCTGTGACCTGAGAGGTCTAGATCTTGTTTCTACAGAGGTGG

CCCTGCAGGACAAGGATAGAACACCTGAGATACCCATCCAGGACTTCAGTCAGTCCTAGG

ACCTGGATCCCCCCACATTGTGTGCCCTGTAGAGCTTCCCATGACAGGCCTCACCATCGG

TTGGGCCTCTTGGGGAATAGGCAGGCATTAGGATGGGCAGAGGGAGCCCAAATCTGGGTA

TGGCGCCCCTTCTTCTGCAGCTGACGCAACGCTCCCTTCTCTCACCCGCTGCAGGCCACA

GAGGAGCCCGTGGTGCCTGTGGAGCCTCTGGAATCCATGCTGAGGCTGGGGCCCAGAGAG

GAGAAACAAATTAGGTTGAGACTCACAAGTTGAGCAGAGCTGGAACTCAGGGTCAAGCCC

TCTGTCTCCTAGTCCGGGATTCCTTCTTCTATTCCTGAACAGGCTGCTTCCCAGTGTCTG

GCCTCTGGGGAAAGCGAACAGGCCCC

**>Pig_predicted**

AGGGCATCCGTACCTCCCCTGCCAGGCAGACACAGTTACGTGGGAGACTCATTACTCCTA

CCTCAATCCCCTCACTGTCCTGGGCCCCCCCTCCTATGTGGTTCTGCCTCCACCTCAGAG

GCCAGGCACCAGGAGACAACCTGGAGCATCAGGATGAACCCTGAAGCTAAGCCTGGACTC

AGGTGGGCTCCCTCTGAGCCCTCAAGCCTCAGGGCCCCAGGACGGTACCTGTGAGCTCCC

CTTTCTGCACAGTGAAGGGAATGACCAGGTTGAAGGGTCTCAGCATGGACTCCATCGACT

CCACAGGTACCACTGGCTCCTCTGTGGCCTGCAGCAGGGGAGAGAGAACAGTCCTTGGCT

GTGGAGCTTGGCCTGAGCCATGATTGACATCAGAGAAAGTGAGGAATGAACAAGATGGTT

AATGGTGACATCAGAGGGCAGGAGAGGCGCGTCAGCTGCAAGAGAAGAGGTGCCATGCCC

AACCTGGGGGGTCTGTCTGCCCAGCCTGACTCCCGCCTGCTCCCCGAGATGGCCCCCAAA

CCGTGCCCCAGCCCTGGCTACT

**>Frog_predicted**

GCTTTTCAAACCTATTTTTATGACTCACTAAAAGTGCTTGTTTTCTGGTTATTTACAAAA

GCAGCACAGGTAAAGGAAAAATAAAGGGCTCTGGCAGGTCAGTCGGACCCTGGACTTTAT

TTTTGCTCTAACTACTGTGCAATTTAAAGTCTGTGAATTTTCAGGAACATGCCTGGGACC

CTCAATTCAGATTGGGGAGGAGACTGTCATCACAGTGGATGCTAAAGCTGCAGGGAAAGG

CAAGGTGACCTGCAAAGTATCTTGATGTAGTGGAAAATCAGGATGGAACTTTTGACATTT

ACTACACAGCTCCTGAGCCTGGCAAATATGTAATCACAATTCGTTTTGGTCCCTTCTGCC

AGGCATGTGACACCATTCTCTTTCTTTTCATCCAATACAGGCCACAGAGGAGCCAGTAGT

TCCTGTAGAGAATTTGGATACCATGCTGAGGCTACTGTGCAAAAAGGGGAAACACAATAA

CATGACACCCACAATAAAAGTTGACTCTGAAAGACACATTGAATCTAGGTGAAGTACGTA

TGCCTTCTGGTAAGACAGCTCGTCCTAATATTACGGATAACAAAGATGGCACTGTCACTG

TGAAATTTGCACCTGTTGAAAATAGACAGTTCCTATCTAGACATGAC

2. **LOCUS OF INTEREST: GATA6/GATA6-AS1**

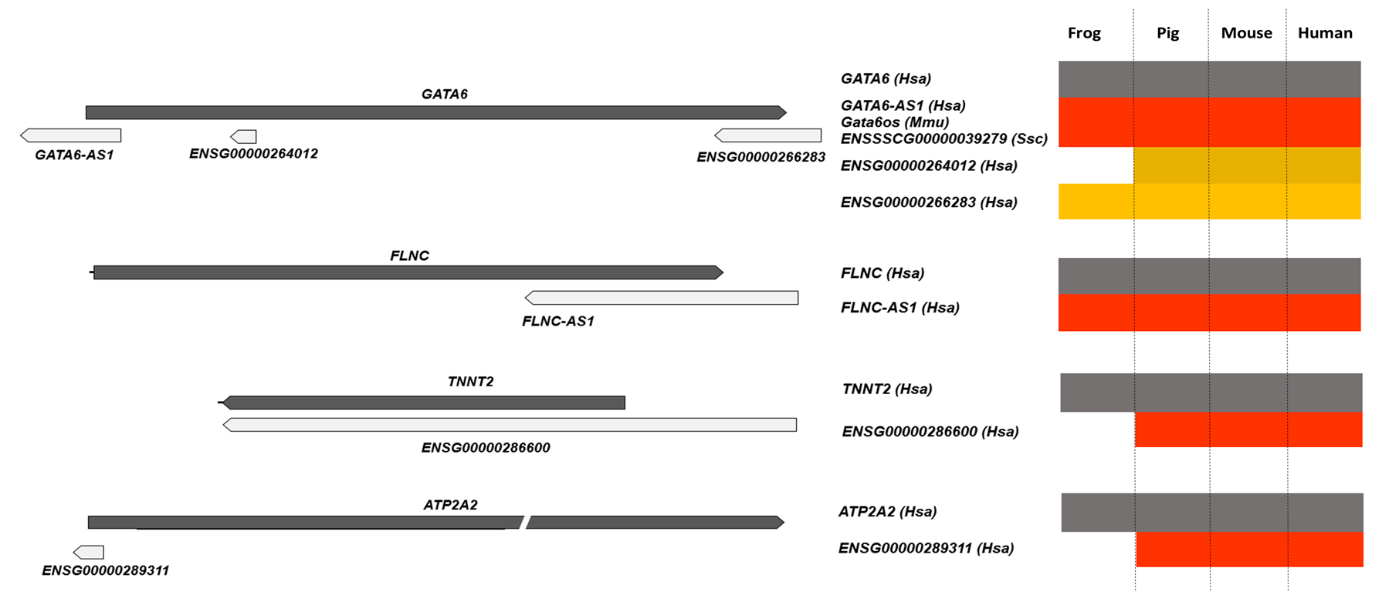

**PREDICTED PRODUCTS**

**>Human_GATA6-AS1**

AAAGGCTGAAGGTAGAAAACTCTGGAATGGAGAGGCTGGCGACTGGAGCAGGATCCAGGG

GCTTTAAATTTTAACCTGAATGCGATCGAATGCGCGACTCCCAGAAGAGATAGCCAGGCG

TCCGCCCGGCCGCTCAACCCCCGCCAGGACTCGGAATGCCTTGCTTTTGGTTCTTTTCCC

TGTTATCTAGGCCACCCAATGACCGACGAAAGATTTTCCCTCCCTGCTTCTCCCGTGGCC

GCTTTAAACCAAATAAATTTAGCTATTAAGGGCCAAGATGTCTGACAAAAGATCGAAAGG

TGTCTTTCCGTGATCGCTTAAAGAAGCGAAGGAGAAGCTTGCGGCTCTCAAATCCCGATA

AAGAAATTCCGCGCCGGCGCCGGCCTCCCTCCCCGCGGCCTGGTTGCTCCCTGACGCCCT

GAGACCCCGACTTCGGCGGCCGCAGACCTCGGACGGGCCCGGCGGGACCTGTGGCTGCCG

CCTGGGGAAGGTGGAGGTCCCCGGGCTGCGGGGCCAGGGAGTGCCAGGCCCTAGGAGACG

GGCGGACAAGGGGGTCCCCGGGAGCTCTGGAGCTCCAGGTGGGAGCTGGGTATCCTAAAA

CGGCCCCCGTGCACTTCGCGGAAAAACTAAGCCAAGCTCAGGGGGCCGAAATTACCACTC

CGTTTTGATTATTTTCTGATACTAAATAATAAAGATAAAACCAAACTGCC

**>Mouse_Gata6os**

AGCACTGGGGGGCGGGTGGAGGCTGGTCCGGAGTATCCCGGGAACCTCGGAGTCCCTACA

ATCTTCTAAAGCGAAGGACCGGACACCCTCGGAACCGATCACAGCCCGGGAAAGGCTGGA

GAGCTGGAGGCCCAAGGGCCCGCGAAGGGAGAAGGGAGAGGAGGGGGCAAACAAAGGAGG

AAACAACCGAACCTCGAGCGATGTGCGAGAAGAACGGCCGCGACCGAGGGTGCTCCGGGG

CGCGGAGAGAGGCGGAGGGCGCCGGCGGGAGAGCGAGAGTCCGGGGTCTGCTAGTCGCTG

CTGGTGAATAAAAAGGAGAGAGGAGGCGCCTCTTACTGCTCTGCCGGAAAACTGCAGCCT

GCGCTCCTGATTGGACTCACCGAGCCCTAAACAAACAGCGCTAGCCGAAGGGTGCCAGGC

TGTGGGTCGGAACTGCTCTCTACAAGCTGCTGGGTGCGGAGCTCACAAAAGGGATTTCTT

TCGACAAGCTGGACATTGGAAGAAGGCTGCGAGAAGGGAATTGAACATGAGAAACACTCT

TGTGGGAGTGGATTCTTACAGGAACACCCCAGTCACTGTGGCTTCAGCGTGGCTGCATTT

ATGAAAGGCTGGAAGTAGAAAACTCTGGAATGGAGAGACTAGCAGCTGGAACAGGATTCA

AAGGCTTTAAATTTTAACCCGAGTTCGATCGAAAGTTACTACCACAAGAGACTGGCACGG

GCCGGCCCAATTCCCACCACGGCTTCCAGCGGCTTGCTTTGGGTTGTCCGTAGGCGTCGA

AGACTAGAGAACACACTGACCCTGGCGCCTCTGGGACAGAGGTGAGGAGTTCGACCCAAA

CGAACAGATTCGACAACATTCGGACTCTAGAGGCTGGGGAAGAAACCCATAATTTTACCG

TCCTCATTCCCAATCATTTTAGTAGTGCCTCCAACCACAAGAAAACCGATAGATCATTGT

TTTTCCTTTCAGGGCTCAAAATTCACCTTTTAAACTTTTTTTAAAGACTAGCTTTCCGAA

AGAGGAATAGGTTAGAGAAATATTTTTTTTTCCATCTCAGTTAATGTAAAAATAAAACCT

GTCCGGTTGACTAAGACAAAAA

**>Pig_ENSSSCG00000039279**

AGGAGAGAAAAGAGGGTGGGGAGGGGAAGGCGGGCGGCGGGGGAGGGGAGCGCGCTGCTA

GGGCAGACAATGAGTCCCGCGCAGCCTGCGGGTGTGGGCTGAGCTGCGAGCGCGGTCCTA

GGACTAAGAGCAAGTCCACCCAGGGGGCGCCGATCCCGGGGACGCCGGGGGAGGGGCAGA

GGTGGCAGAACGCCTGACCTGGCTGGCGACCCACACCCATTCCTCCTCCCCCAGTTACAC

ACATACTCAAGGGGGGGAGGCGCTGGCAGGGTTGACTATGCATAGTCCAACGCTGTCTCC

GCAGCGGGAGAGACGCGCCATCTCGGGGGTCAGCGAGCCCTCAGAAAGTGAGGGGAAAGT

GCCTTCCTCCGCTGGTCTCCCCCACTGCTGCCTCCACATTAAATGAACTTTTTGCCCCGT

AGTTGTGTATGTTTCCCGACCTCTTCTCGAGACCTTGGGGCTCAGGTCCTGGTGGCCGTC

GGGGGTCTAGTCTCCTGTGATGTAGGAAAGAAGGAACTAGAGCGCCTAGCTCCCTATCTC

TGCTCCGCTTTTTACAAACGGTACTAAATTCCTGGGAAGTTTCGGGAGGTACACGTCGGC

ACTCCCCAGCTAATCTCTGAGAACTTTTAATTCGGGGCCCTGGTGTACGTAAAGGTTAGT

CCACGAGGTGCGCAGAAGGGACAAGGGCTACTAGGCCAGTCGCGCTCTCCGAGTTCTGGT

GTCCTGTTTGCTCTGAGCTTTCCCTGGAGAACTTCAGGTAAGTCACATATAAGTCGTTGG

AAAGAGGAGGGGGGTCACCTTCCCTGGTTCCAGGGCTAAATACCACTTCCCCACCCCCAC

CCTCACATACACACCCCTTCTTCGCTCCCTCCAAATGGTTATCACAACTGTGTACAAACT

GATACGGTCCTTCGCCGGCAAAGGGACTGGAAATGAACGTCGCGGGGTTATCAGCGCCGA

TCTGTCAGCAGGGAGTCGCGCGGCCCGCAGGTTCGTCCTCGGCAGGAGCCGTGATGTATG

TCTCGCTCCCTTACGCTCACAAAAAGGATTTCTTCCGACAGCTGTGACCCCAGAAGGCTG

TCGAAAATTTTAACTAGGGCAGGAGGAGTTGTTTTGTAGGAATGGATTTTTACACTTTTT

TTCCAGGTAAATCGAAGTAAATGATTTTCTAAAGGTCAGTCTCCCTGAAAGCCTTTGGGC

TGTGTGTGTGTGTGTGTGTGTGTGTGTGCGCGCGCGCGCGCGTGCGCGCGCGTGTGTGTG

TGTGTGTGTGTGTGTGTGTGTGTGTGTTTCCAGTTGGGGAAGGGAGAGTTGCATAGAGTT

GAACCCGGGTAGGTAGGGACATAATAAAATTTAAGAGTTTGGAGAGATTGTGAGAAGTCA

TGGCTAAATGTTTCCTTTCTATGTAACTGTGACTTGTGAGCCTGTGTATCCTGGGGGACA

AGGGGACAGGGCCAGGCAGCGCGGGGGAGGAAGGTGCAGAATCGAGGAAGTCCACGATTC

TCCTTTTCTTTTTTTCATTCTTCGTTGTAAATAGCTGTTAAACACTTCTGAATTTTAAAT

TGCTACCGAGAAGGTCCATAAACCGTTCTCATCCAATTTAGTCACTTTTATTACATTCCT

CTTCGCCTCCCCCAATCACTGCGGCGGCTCCCAGTTTCAGCGTGGCCGTAATTCGGAAAG

ACTGAAGGTAGAAAACTCTGGAACGGAGATTGGCGACTGGAGCAGGATCCAGGGACTTTA

AATTTTAACCTGAATTCGATTGAATGCATCTACCAGAAGAGATAGTCCGGCGTCCGCCGT

CCACTCAACTCCGACCAGGGCTTCCAATGCCTTTCTTTTGGTTTCTACCGAGATGCTGTC

TAAGGAAGACAAGTTAGCAAATAAATTGGCTAAGTAAGAGTCCAGCCTGTGAACGGCCTC

CAAGCCTGAGCCCAGAGAAGACCGAGCCCCGGCACGAACCTGATTCCTGCTGCGTGTGTG

TTTTTTTATATTTGCCAAACACGAGGTTGTCCTTGTGAATTTCCCTGCACAATAGCTGAC

ACCCGGGAGAGGGTGCGCGGGCCCCGGCGCCGCTGGGACAGAGACTGGCTTTCCAACCAG

AGGAGAGAATGGATAGTGAAATTTTTTTCTTGTCAGTTAACTTCAAAATAAAACAAAACT

GTTTGACCCGTGGAGTTTATTCAGACGGAAAGCCAGGTTGCCTGACTCGTCCAGTTTGCA

ATTTTCCCAGCGCATCGGCTTTGGCCCCGGAGACAACACCCCCAGGGAGAGGCACATTTT

CAAAGCCACCCTTCACCGGCCCCAGGCCCGGGGCTCTGATCTCCCTGAGCGCCTCAGTTG

GTCCCGGAAAGCGAGGAGAAGAGGGGGTGCAGAGGAAGAAGTGCACCCAAAGAGCTGGCG

AAGGCCCCCATAGTGTTCCAGGTGGAAAGCTTAGAAGTTTTCTTCGTGTTTCCTCCCGTT

TCTTTTCTTCCTCCTCTTTTCCTTCCGCCCCTCTCTTCTTCGGTGATTTGCTAATATGGA

ACGAAATCGTTTGTCTCGGGACTATTTAGACCCAAGAAGTGGCCCCTCCGGAGCGAGCCT

GCCTCCCCAAGCGGCCCTTCTCACCTTCTGAAGGGGGCTGGCTCCCGGGCGCGGCCTGTC

TCCAGGCTGATTTATTTGGGCCGCGCGCGCCACACCGCGCTTCGCGATTCGAATCCAGCC

CGAACCCAGCCGCCAATAGGGCAGGAGGCGGGGGCTTGGCGGCCAAACGGGGACGGGTGT

TTGGGGACCAAGTTGTACGCCGCTTCCGGCGGAGCAGCGAGACTCGGGCGAAGACCCGTG

TCATGATCCGCCTTTCTTCCCCAAACCTAATCTCATCCTGTTGTCCTTTCACCAAGCTCC

TGGGTGCTCCAGCGACGTTGCAGGAGGAAAAAGAATACAATTACATACCGTTTCTGACTT

AAAATATAGTTTTAACTT

**>Frog_predicted**

GACAGAGTGAGCACGACCTAGTTACGGAGAGGAGGGCGCTCCGGGCATCTTCAGGGTGCA

GCTGATTTTGGGGGGGGGGGGGGCACAGTGGGCGTATACCAGCACAACTAGTCAGGCAGC

CTTCGGCTCTCTGGGGTCTCGGCAGCACCCGGAGAGCCGCTGGTTGTGTGTCTCTCTATG

GATACGTTGCTCACTGAAGTTAAACGACTAAAGTTTTTCATATCCCAGCGCCCCCACACC

TGGATGGACCTGAATGAGGACAGCTGGTCCGGCGGCACAAGGCTGAGTCCCGCGAATTCC

TTCTTCCCCTGTCCGATCAAACAGGCAGAGGGGAGGCCGGCTCCCTCGTATCACCCGCTC

GGCCTCTCCCACCCCGAGGAGGGAGACTCGCTACAGCAGCGCCCTGCCCGCCCAAGGACC

GCTCGGCTACGATCACTCTCAGGGCACTTTCATGCACTCCGCTGCCAGCTCGCCGGTCTA

TGTGCCCACTAGCAGAGTTGGCTCCATGCTCACAAGCATCTCCTACTTGCAGGGCACCGG

GGCATCGCAGGGCGCCCATTCAGTGAACAATCACTGGTCGCAGGCAACCTCCGAGGGTTC

ATCATTCAGCAGTAGTAGCCCGCATACCTCTAGCCGGTACCACTACTCCCCCAGCCCGCC

CATGGCCAATGGATCTACCCGGGATACAGGTTAAGCTCACCTCCGCTTGGCCTGCTGGAC

CTTTTGACAACACCATGCTTCACAGTCTGCAAAGTAGCATATATTTGGAAAACCCTAATA

TGTTGGTAATTTGGGAATTTTATAAATCACAGTTTTGTCATGAAACTTTGCTAAAAACAA

GATAACTTGTTCAAATGGAACCA

3. **LOCUS OF INTEREST: ATP2A2/ATP2-AS1**

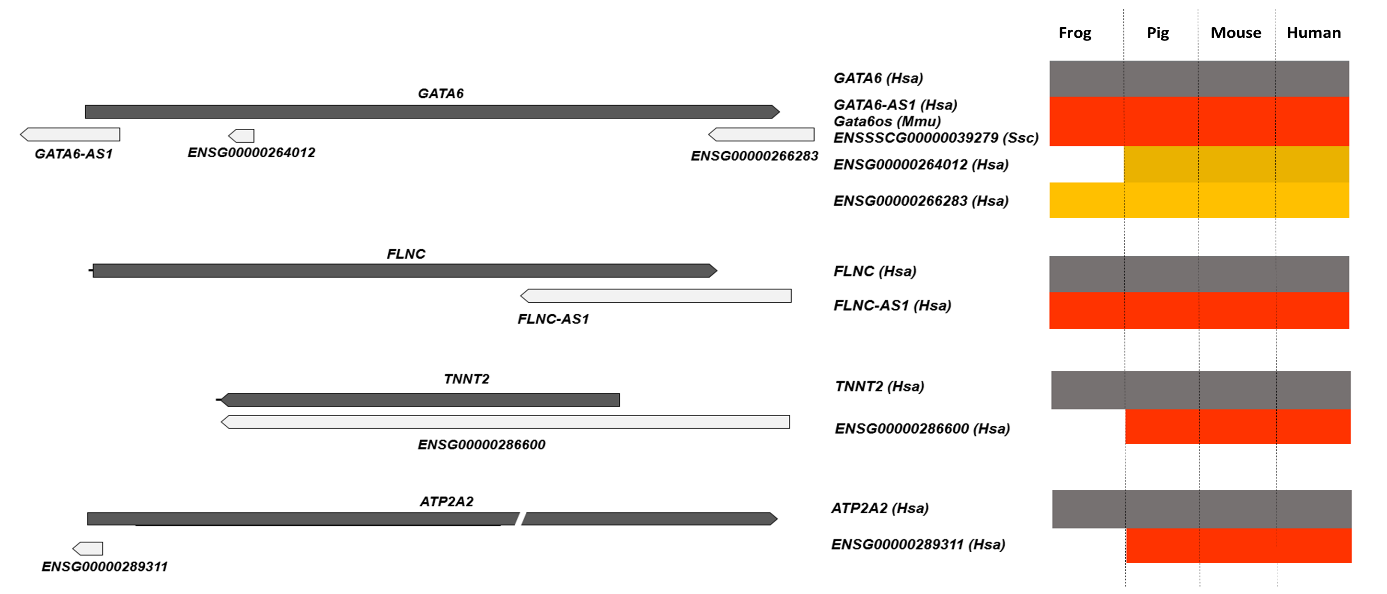
 Frog	pig	mouse human

**PREDICTED PRODUCTS**

**>Human_ATP2A2-AS1**

ACAGTTCACGCCGCTGGAGCTGCCACACAGCAGCTAGACTTGGGCTAGGAGGTGCTTCGC

TGTGGCTTTCAGATGAAGGCCATTTCATAGACTCACTTCCCTCGACAGGAGAAAAGGGGA

CTTGAATCCAAAGCTGCTCTTAAACCACAGATGATTTGCTCTGCTCGGGAGGACAAAGCG

CAGGTCACAGCAAGATAAAGGGAATTTTGCCAATATAAAGAAGCTACATTACTTAACAAG

GGGATGGGAAATTGCATGGAATTCATGAACAAAACATGAACATATTCTCTTCACAAAGAA

GAGCAGCTTTCTTCACGGAGTAGGTTGCAGAATGAAGCAATTAGTCCACTTTTGTGTTTA

ACAATAACGCACCATCCGCATGAGCAAAAGCAGAGCAGAAAAATCACCTCAGTTGTCTCA

AGTGGCTGGGCTTCAGGACTTAAGATAAATTCCTTCAGCAGATGCAAATTACAGTGCATC

ACCAGCTGAAAAAATCAGGCACGTCATGTCATTTAATTATTTCCAAGGAGTGGACGTTGG

CATGTAGTTTTGATTGATACTTGGAAAAGAACCAACAAAACCCTACACGAAACAAACAAA

CAAAAAATTATACTATATGGCAACGCACTGAGAGAAGCCAGGTGGAAACTGGTTAGGTCA

CAGGATCAGAGCCGCTTCAAGAGGGGCACGTGAGTGCACGGTGGGAGTGTGGTCATGGAG

GGCTGGCTGTTGTTGCCCTCTGAGACCAGCTCTGCACCATTTCGTACCAATTCTTCTCCC

TTACCTCCTCTGTCTCTTTCAGGACCGATAGAAGCACTGCAGGCAGGAGAAGGATGGTTT

ACCTTTCTCAGCCTGGAAATTAGCTTAGGCATTTGATGTTGAATATTCTGTGCCACTGTC

ACTGCTTGGGGTTGTAAGCACCTCTGGTGCAATTATGAGCTTCTGGAGGAGTTCCTGACC

TCAGCCTTTGTATGGAATGCTCTTCCCACAGATCTTCTTCACTTGTGTCGTTCTCTTTCC

ATTCAGATCGCAGCTTACATGTCCCCTCCTCAGAGATGTCTTTGCTGTCTAAAGTGGCTA

CCCAGCCACAGTGGCTCACGTCTGTAATCCCAGCATTTTGGGAGGCTGAGGCAGGAGGAT

CCCTTGAGCCCAGGAGTTTGAGACCAGCCTGGGCAATATAGCGAGACTCCATCTCTACTA

AAAAAAAAAAAGAAAAAAATTAGCTGGGCGTGGTGGCCTGAACCTTTAGTCCCAGCTACT

CAGGAGGCTAAAGAGAGAGGATCGCTTGAACCCAGGAGGTCAAGACTGCAGTGATCTGTG

ATCACGCCACTGCACTCCAGCCTGAGCAACAACAGAGTGAGACCTTGTCTCAAAAAAAAA

ATTAAATAATAAATAAATTAAAAAGTGGCTACTCTCGGCAATTCACCTTC

**>Mouse_Atp2a2-as_predicted**

GCTTGGCTTTGGCTTCCAAATGATGGCTGCTATATAAGACTTCCACTTCCCCCAATGCGA

GAGAAGGAGCCTTCAATCCCAAATCTATTCTTAAACCACAGATCATCTGTTGTTTCTGTG

AGGTCTGTACAGAGCCTAGCATGCAAGGTAATTTTGCCAATATAAAGAGGCTATACTACT

TAACAGGGAGGAGGCCCTTTGCATTGAATTTGCAAATATCTTCACAAAATGCGGCTTCAT

GAAGGGCATAGATTGCAAAATAAAGCATTTCATCTTCTTGAGTGTTTAGCAACGCCACCC

CCCCCCCCCAATCACTTGAGACGTCCCACAAGGCTCCGCCTCAAGACTTAAGATAGATTC

CTTCAGCAGATGCAAATCATGGTGCTTTACCCGACTGAAAAAAAAAATCAAGTACGTCAC

TTCATTTAATTATTTCCAAGGAGTGGAGGCTGGCACGCAGTTCTGATTGATACTTGGAAC

AGGACCAACAGAACCCTACACGAAACAAACAAACAAAAATTCTAGTGTATAGCAATGCAC

TCCCTAGAAGCAAGGTGGAAACTGGTTAGGTCACAGCCTCTTCTGGAGGATCAGGTGACT

GCCTGCGAGGTGGGGGGTGAGAGGGTGGGAGGGATGGAGAGCTGGTGGGTCTCCCTCCTA

GCTGGATTCCAGTTCTGTCGTACCTACAGCAAGGGGCACTGAGAGTAGGAAGGTCATGAC

CTCTCTTAGTTTGTCTGTGATGCTCACTGGGTTTGGCTTTTGTTCTGTAGTCCCCTCCTT

CAGGGGCTGGAGGTGCACCCTACAACACGGGTTCCCAGTTTTTCCCTCAGGCCATCTTCC

TCCTTGTCCGGTCCCCTCCTGAAAGATGTCTTTATTATCTAAAGCAGCTCCCCTCCCCAG

GATGCCTACTATCACACTCCTAGCATCTTGTAAGGGCATCCAGACTTGAGAATGAAGAGT

GCCTACAGTTTAAATGCATGAAACCAATGAGAGCCCCACCCTCCTCAACCCATCAGTTAC

TTGGGAAGGGCTCTGCCAGCACTAAGGCCTCAGCAAAAGAAGAATGGCTGTGAAGCTTAG

GTTTAAACCAAAAAGAAGGGCTGGAGAGATGGCTCAGCGGTTAAGAGCACTGACTGCTCC

TCTGAAGGTCCTGAGTTCAAGTCCCAGGAACCACATGGTGGCTCACAACCACGAATAAAT

AAATAAATCTTAAACAAAACAAAAA

**>Pig_ATP2A2_AS1_predicted**

ACAGATTACGCTACTGTAACAAATAGCGAGAGTTGTGCAAGAAGACACTTCGCTTGGTTT

TTAAGATGAAGGCTATCTTCTAGACTTTCACTTTTCCCGATAGGAGAAAAAGGCACTTGA

ATCCAGAATCTGTTCTTAAACCACAGATAATTTGCTCCTGAGGACTATAGAAAACCTACC

TAACTGCACAATAAAGGGAATTTTGCCAATATAAATAAGTTAGACTACTTAATAGAAAGG

GACCTTTGCTTTGATTTCATGAACAAAAGGTGAACACATTCGCATCACAAAGAGCAGTTT

TCTTAAGGGACTGTGTCGCAGAATGAAGCATTTAGTCCACTTTTGTGTTTGGCCATGGTG

TACCATACACAGGAGCCAAAGCAGAGCAAAAATCACCATAGTTGTCCCAGATGGCTCAGT

CTCAGTACTTAAGATAAAGTCCTTTAACAGATGCAAATCACTTTGCATTACCCCACTGAA

AGAAAAATCAGGCACCTCATTTCATTTAATTATTTCCAAGGAGCGGAGGCTAGCATGTAG

TTTTGATACTTGGAAAAGGACCAACAAAACCCTACACTAAACAGAAATTCTAGTGTATGG

CAATGCGCTAAAGAGAAGCAAGGTGGAAACTGGTTAGGTCACAGAATAGGAGCCTCTTCC

AGGAGGGACATGTGACTGGATGGTGGGGGTGTGGTAATGGAGGGCTGGTGGTTCTTTTCC

CACTCAGACCGCTTCACCACTTCTGATTCTGACCAATTCTTATGCCTCACCCCAGCAAGG

ATTGTTACCTTACTCAGCCTAGAAATTACCTTGAGCATTTTACTTTGGACCTCTGTTCCA

TGGCTCCGTCTCTGCCTGGGGTTGTAAGCACTTCTATGATTATAATCTTCTTGAGAGTTT

TCTGATCTGGGCTCTTGTACTCAATGCTCTTCCCTAGATCTTCATGCTCCTACTTCCATC

TTATTCAGATCTCAGCTTATATGTTACCTCCTCAGAGATGTCTTGACTGACTACTTAAAG

GGTTATCCTCCCCAACAAACCTTGCTGTCATATTATTTTTTTCTTCAAAGCATTTTGGCT

CACCATACCAGGATGTCAGCTCCATGAGAACAGAAGCTTGTCTAGCTTGTCCATCTTCAG

GATGCAGAAAGCTCCTGGTATTCATTCATCATTTACTGAATGCCTACAGTTGTTTGTTTG

TTTTATGGCCATCCTTGGCATATGGAAGTTCCTGGGCCAGGGATTGAATTCTAGCTGAAG

CTTCGACCTACACCACTGCTATAGCAACACAAAACCCTTTAACCCATTGCAGTCACACCT

CCACAGTGACCTGAGCTATTGCAATCAGATTCTCAACCCACTGCACCACAGTGTGAACTC

CACCTCCTGTTTCAGTGAATGAAAGAATCTTACCATCCCTGATTTATAAGGGACTTTAAA

AGGGCCTGGTCATGCAAGGGCCTCAGCAAA

4. **LOCUS OF INTEREST: TNNT2/TNNT2-OT1**

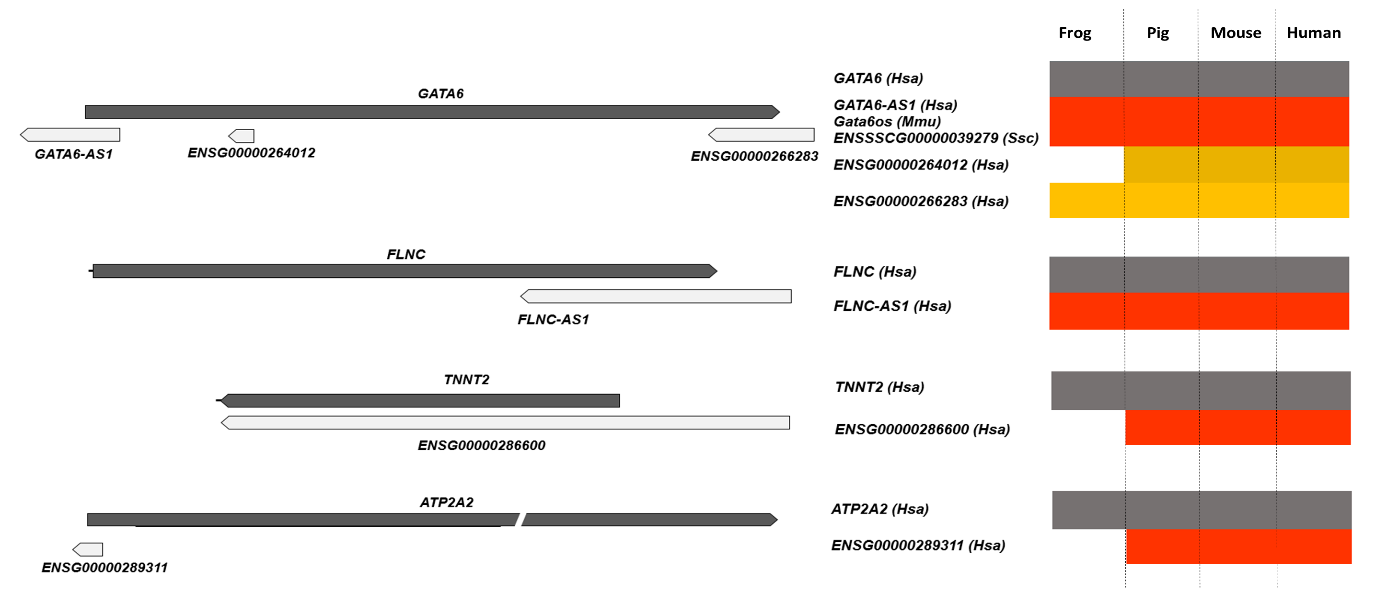
 Frog	pig	mouse human

**PREDICTED PRODUCTS**

**>Human_TNNT2-OT1**

GAGTCCCCGCTGAGACTGAGCAGACGCCTCCAGGATCTGTCGGCAGCTGCTGTTCTGAGG

GAGAGCAGAGACCATGTCTGACATAGAAGAGGTGGTGGAAGAGTACGAGGAGGAGGAGCA

GGAAGAAGCAGCTGTTGAAGAGCAGGAGGAGGCAGCGGAAGAGGATGCTGAAGCAGAGGC

TGAGACCGAGGAGACCAGGGCAGAAGAAGATGAAGAAGAAGAGGAAGCAAAGGAGGCTGA

AGAATGGCCCAATGGAGGAGTCCAAACCAAAGCCCAGGTCGTTCATGCCCAACTTGGTGC

CTCCCAAGATCCCCGATGGAGAGAGAGTGGACTTTGATGACATCCACCGGAAGCGCATGG

AGAAGGACCTGAATGAGTTGCAGGCGCTGATCGAGGCTCACTTTGAGAACAGGAAGAAAG

AGGAGGAGGAGCTCGTTTCTCTCAAAGACAGGATCGAGAGACGTCGGGCAGAGCGGGCCG

AGCAGCAGCGCATCCGGAATGAGCGGGAGAAGGAGCGGCAGAACCGCCTGGCTGAAGAGA

GGGCTCGACGAGAGGAGGAGGAGAACAGGAGGAAGGCTGAGGATGAGGCCCGGAAGAAGA

AGGCTTTGTCCAACATGATGCATTTTGGGGGTTACATCCAGAAGCAGGCCCAGACAGAGC

GGAAAAGTGGGAAGAGGCAGACTGAGCGGGAAAAGAAGAAGAAGATTCTGGCTGAGAGGA

GGAAGGTGCTGGCCATTGACCACCTGAATGAAGATCAGCTGAGGGAGAAGGCCAAGGAGC

TGTGGCAGAGCATCTATAACTTGGAGGCAGAGAAGTTCGACCTGCAGGAGAAGTTCAAGC

AGCAGAAATATGAGGCCACCTGGGACCTGAGCCAGTCAGCTCCAGCGTTGCTCTTTGTCC

TTCCCACTTTTCTTGCAGATCAATGTTCTCCGAAACAGGATCAACGATAACCAGAAAGTC

TCCAAGACCCGCGGGAAGGCTAAAGTCACCGGGCGCTGGAAATAGAGCCTGGCCTCCTTC

ACCAAAGATCTGCTCCTCGCTCGCACCTGCCTCCGGCCTGCACTCCCCCAGTTCCCGGGC

CCTCCTGGGCACCCCAGGCAGCTCCTGTTTGGAAATGGGGAGCTGGCCTAGGTGGGAGCC

ACCACTCCTGCCTGCCCCCACACCCACTCCACACCAGTAATAAAAAGCCACCACACA

**>Mouse_Tnnt2-ot_predicted**

GAGAGCCCCCTGGGAGGGAGCACACCTTGTTCGAGGAGGAGTTGTGGGAGGAAGGAGTCT

GGAGAAGGGTCTGGTAAGGCACCTAGGCCCCTTCCTTGACCCTGCCTCATTTTCTCGGGG

GCCGGGATCCAGAAGCTAATGATGTGTCATGCTGCTAAAGCAATAGGGGAGGAGGGAGGG

TCAGGGGACGGGGAGGAACCATCTCTAGCCAAGTGTCCCTGCTCACCATCCCCTTCAGAC

AGAGCGGAAGAGTGGGAAGAGACAGACAGAGAGAGAGAAGAAGAAGAAGATCCTGGCAGA

GAGGAGGAAGGCGCTGGCCATCGACCACCTGAATGAAGACCAACTGAGGGTGAAGGCTGA

GGAGCTGTGGCAGAGTATTCACAACCTGGAGGCTGAGAAGTTCGACCTGCAGGAAAAGTT

CAAGCAGCAGAAATACGAAGTCGACACTGAGCCCCCTCCCTCCTCCCAGGCCTCCTTGAA

TGAGCCCCAACTTTATAATTTATCTTCCCTGCTTCCCCTCCCCCGCCCCCAGATCAACGT

TCTGCGAAACCGGATCAATGACAACCAGAAAGTCTCCAAAACTCGTGGGAAGGCCAAAGT

CACCGGGCGTTGGAAATAGATGAAACTGTTCTCGTCAAAGCTGTGCCCCCTGCTTGTGTC

CTTGCCCCGTGCATCCCAGCTCTGGGTCCTCCTTGGCACCCAATGCAGACTCCTGTTTGG

ATAGTGGGAGCTGGCTTAGCTAGAAGCCAGTACTCTGCCTGACCCCTATGCCCGCACTAT

GCCAGCAATAAAAAGCCAACACAC

**>Pig_TNNT2-OT_predicted**

CAGGCTGTTTATGCCCAACCTGGTGCCACCCAAGATCCCCGATGGAGAGAGAGTGGACTT

CGATGACATCCACCGGAAGCGCATGGAAAGGACCTGAACGAGCTGCAGACGCTGATCGAG

GCGCATTTCGAGAACCGCAAGAAGGAGAGGAGGAGCTGGTCTCTCTCAAAGACAGGATCG

AGAAGCGCCGGGCGGAGCGGGCGGAGCAGCAGCGCATCCGGACAGAGCGGGAGAAGGAGC

GGCAGACCCGCCTGGCTGAGGAGAGGGCCCGCCGGGAGGAGGAGGAGAACCGGAGGAAGG

CTGAGGACGAGGCCCGGAAGAAGAAGGCTCTGTCCAACATGATGCATTTTGGAGGCTACA

TCCAGAAGGTGGCCCAGACGGAGCGTAAGAGTGGAAAGAGGCAGACGGAGCGGGAGAAGA

AGAAGAAGATTCTGGCCGAGAGGAGGAAGGTGCTGGCCATCGACCACCTGAACGAAGACC

AGCTAAGGGAGAAGGCCAAGGAGCTGTGGCAAAGCATCTACAACCTGGAGGCGGAGAAGT

TTGACCTGCAGGAGAAGTTCAAGCAGCAGAAATACGAGGCCACCTGGGCCCCGAGGGCAG

CCGATGCCAGTGATGCTCTGTCTTTCCCTGTTTTCTTGCAGATCAACGTTCTCCGAAACA

GGATCAATGATAACCAGAAAGTCTCCAAGACCCGCGGAAAGGCCAAAGTCACTGGACGTT

GGAAGTAGAGGTGGGGAGCTGGCCCAGCTGGAAACCGCCTTCATGCCTGACCCCACACGC

CCGCCCCAACCAGCAATAAAGGACACCACGCA
