## Supplementary material for "An integrative cardio-informatics approach identifies positionally conserved ancestors of human lncRNA genes transcribed across vertebrate hearts": Supplementary File 2.docx

**RNA isolation and reverse transcription on mammalian heart tissues**

50–100 mg of tissue was transferred to 2.0 ml Eppendorf tubes containing 0.5 g ceramic beads (1.44 mm; Peqlab, Erlangen, Germany) and 1 ml TRIzol reagent (Qiagen, Germany). Following homogenization with two centrifugation steps (6.000 rpm, 30 sec) in the Precellys homogenizer (Peqlab, Erlangen, Germany), the suspensions were mixed with chloroform, incubated at room temperature and centrifuged at 12.000 g and 4°C for phase separation. The water phase was transferred to a new Eppendorf tube, mixed and incubated with isopropanol for 10 min at room temperature, and subsequently centrifuged for 20 min at 12.000 g and 4°C. The RNA pellet was washed with 75% ethanol for 5 min at 7.500 g and 4°C, air-dried and dissolved in nuclease-free water (Qiagen, Hilden, Germany). RNA sample purity and concentration were evaluated in a NanoDrop ND-1000 spectrophotometer (NanoDrop, Peqlab, Erlangen, Germany). Reverse transcription was performed on 1.2 µg of total RNA, following the instructions of the GoScript™ Reverse Transcription System (Promega, Madison, Germany).

**SUPPLEMETARY MATERIALS AND METHODS**

**RT-PCR for detection of lncRNA genes in vertebrates**

Predicted lncRNA orthologues were validated for four selected candidate PCGs. The PCR protocols described below were carried out in a Biometra TAdvanced Thermal Cycler (Analytik Jena, Jena, Germany). Each 25-µl reaction mixture contained 1 µl cDNA, 0.15 µl HotStarTaq Plus DNA polymerase (Qiagen, Hilden, Germany), 2 µM of each sense and antisense primer (Table 1), 1 mM dNTPs (Thermo Fisher Scientific), 2.5 µl 10× buffer (Qiagen, Hilden, Germany), and nuclease-free water (Qiagen, Hilden, Germany). The touchdown programme comprised two sections. The first section starts with an initial denaturation step at 95 °C for 2 min, followed by either 20 cycles (for mouse ATP2A2A-S1, mouse GATA6-AS1, mouse TNNT2-AS1, human TNNT2-AS1, human ATP2A2-AS1, pig ATP2A2-AS1, and pig TNNT2-AS1) or 25 cycles (frog GATA6-AS1, human FLNC-AS1, pig ATP2A2-AS1, pig FLNC-AS1, and pig GATA6-AS1) of denaturation at 94°C for 30 s; annealing at 65°C for 30 s, decreasing by 1°C per cycle; and elongation at 72°C for 30 s. The second section continues with 24 cycles of denaturation at 94°C for 30 s; annealing at 55°C for 30 s; elongation at 72°C for 30 s; and a final elongation step at 72°C for 5 min. If amplification failed (as in case of frog GATA6-AS1), the HotStarTaq Plus polymerase was substituted with the high-fidelity DNA polymerase Platinum Taq (Invitrogen/Thermo Fisher Scientific) along with 10× high-fidelity PCR buffer (Invitrogen/Thermo Fisher Scientific). The respective 25-µl reaction mixture contained 1 µl cDNA, 0.1 µl Platinum Taq (Invitrogen/Thermo Fisher Scientific), 0.2 µM of each sense and antisense primer, 0.2 mM dNTPs (Thermo Fisher Scientific), 2.5 µl 10× buffer (Invitrogen/Thermo Fisher Scientific), 0.2 mM MgSO4, and nuclease-free water (Qiagen, Hilden, Germany). The high-fidelity PCR program consisted of 35 cycles of denaturation at 94°C for 30 s, annealing at 60°C for 30 s, elongation at 68°C for 2 min and a final elongation step at 68°C for 10 min.

For PCR products that failed to amplify with the methods described above (as in case of mouse FLNC-AS1 and frog FLNC-AS1), a nested PCR approach was used. Two sets of primers (Table 1) were employed in a nested PCR approach to enhance amplification success. In the first round, a relatively distal primer pair and HotStarTaq Plus DNA polymerase was used to generate an initial amplicon, which was subsequently purified (ExoSAP-IT Express PCR Product Cleanup; Applied Biosystems). One microliter of this purified product then served as the template for a second PCR using a pair of more closely spaced primers. For both PCR steps the temperature protocol consisted of 35 cycles of denaturation at 94°C for 30 s, annealing at 58°C for 120 s, and elongation at 72°C for 120 s, followed by a final elongation at 72°C for 6 min. A summary of the PCR conditions used for amplification of individual lncRNAs is provided in the following Table.

**Table 1**: Primer sequences and conditions for the PCR-based validation of predicted lncRNAs

| **Gene product** | **Sense and antisense primers (5‘→3‘)** | **Predicted amplicon length [bp]** | **DNA polymerase** | **PCR strategy** | **Cycle number** |
| --- | --- | --- | --- | --- | --- |
| *Homo sapiens:* |  |  |  |  |  |
| *TNNT2-AS1* | AGGAGAACAGGAGGAAGGCT,  CCTGTTTGGAAATGGGGAGC | 565 | HotStarTaq Plus | Touchdown | 20 + 24 |
| *GATA6-AS1* | GAAGAGATAGCCAGGCGTCC,  GTGCACTTCGCGGAAAAACT | 525 | HotStarTaq Plus | Touchdown | 25 + 24 |
| *ATP2A2-AS1* | CTTGGGCTAGGAGGTGCTTC,  TTCTCTCAGTGCGTTGCCAT | 598 | HotStarTaq Plus | Touchdown | 20 + 24 |
| *FLNC-AS1* | CTGTTAGACGCCGAGACCC,  TCTGCCCAGCCTAACTCTTG | 501 | HotStarTaq Plus | Touchdown | 25 + 24 |
| *Mus musculus:* |  |  |  |  |  |
| *TNNT2-AS1* | CAGAGCGGAAGAGTGGGAAG,  CCCCTATGCCCGCACTATG | 542 | HotStarTaq Plus | Touchdown | 20 + 24 |
| *GATA6-AS1* | GGACTCACCGAGCCCTAAAC,  GGACTCTAGAGGCTGGGGAA | 509 | HotStarTaq Plus | Touchdown | 20 + 24 |
| *ATP2A2-AS1* | CAATGCGAGAGAAGGAGCCT,  CTGGAGGATCAGGTGACTGC | 550 | HotStarTaq Plus | Touchdown | 20 + 24 |
| *FLNC-AS1* | TATGGTTGCCGTCGTACTTG  AATGCCTGCCTATTCCCCAA | 350 | HotStarTaq Plus | Nested,  1^st^ PCR | 35 |
|  | TATGGTTGCCGTCGTACTTG  CCCCAACATCACTGACAACA | 101 | HotStarTaq Plus | Nested,  2^nd^ PCR | 35 |
| *Sus scrofa:* |  |  |  |  |  |
| *TNNT2-AS1* | TGGAAAGGACCTGAACGAGC,  AGCAGAAATACGAGGCCACC | 501 | HotStarTaq Plus | Touchdown | 20 + 24 |
| *GATA6-AS1* | GGGGAGGAAGGTGCAGAATC,  CCTGCTGCGTGTGTGTTTTT | 522 | HotStarTaq Plus | Touchdown | 25 + 24 |
| *ATP2A2-AS1* | AGGGACCTTTGCTTTGATTTCA,  GACCGCTTCACCACTTCTGA | 509 | HotStarTaq Plus | Touchdown | 20 + 24 |
| *FLNC-AS1* | GGCATCCGTACCTCCCCT,  TCTGCCCAGCCTGACTCC | 511 | HotStarTaq Plus | Touchdown | 25 + 24 |
| *Xenopus tropicalis:* |  |  |  |  |  |
| *GATA6-AS1* | TAGTTACGGAGAGGAGGGCG,  GCGCCCATTCAGTGAACAAT | 553 | Platinum | Standard | 35 |
| *FLNC-AS1* | AGAAGGGAGGGAAGTCACAC,  GCAAAAATAAAGTCCAGGGTCC | 516 | HotStarTaq Plus | Nested | 35 |
|  | AGAAGGGAGGGAAGTCACAC,  GTATGTGCTCGCCACCAAAA | 118 | HotStarTaq Plus | Nested | 35 |
