## Supplementary material for "An integrative cardio-informatics approach identifies positionally conserved ancestors of human lncRNA genes transcribed across vertebrate hearts": Supplementary Table S3.docx

**Supplementary Table S3: Microsynteny evaluation of the 109 CMI/lncRNA loci**

| CMI | Neighboring lncRNA | Syntenology conservation (with lncRNA genes in the respective region, if annotated) | | |
| --- | --- | --- | --- | --- |
|  | Human | Mouse | Pig | Frog |
| *CACNA1C* | ENSG00000278255: novel (AS) | Y | Y | N |
| CALR | ENSG00000267458: novel (AS) | Y | Y | Y |
| *CAV1* | ENSG00000235427: novel (AS) | ENSMUSG00000141998 | Y | Y |
| *CHD2* | ENSG00000272888: CHASERR (AS) | ENSMUSG00000108563 | Y | Y |
| *CXCR4* | ENSG00000289974 | Y | ENSSSCG00000061480 | N |
| *DAAM1* | predicted (AS) | ENSMUSG00000127434  ENSMUSG00000127473  ENSMUSG00000127518 | Y | Y |
| *DSG2* | ENSG00000264859: DSG2-AS1 | Y | Y | Y |
| *DSP* | ENSG00000261189: DSP-AS1 | Y | Y | N |
| *FBN1* | ENSG00000259705: FBN1-DT | Y | Y | N |
| *FLNA* | ENSG00000285018: novel (AS) | Y | Y | Y |
| *FLNC* | ENSG00000242902: FLNC-AS1 | Y | Y | Y |
| *GALNT1* | ENSG00000266965: novel | Y | Y | N |
| *GJA5* | ENSG00000274415: novel (AS) | Y | Y | Y |
| *JAG1* | ENSG00000224961: LINC01752  ENSG00000270792: novel (AS) | Y | Y | Y |
|  |  | ENSMUSG00000128110 | Y | N |
| *NEXN* | ENSG00000235927: NEXN-AS1 | Y | Y | Y |
| *NIPBL* | ENSG00000285967: NIPBL-DT | ENSMUSG00000143652 | ENSSSCG00000061094 | N |
| *RBM20* | ENSG00000289954: novel | Y | ENSSSCG00000056393 | N |
| *RPSA* | ENSG00000286781: novel (AS) | ENSMUSG00000111028 | Y | N |
| *SOS1* | ENSG00000229692: SOS1-IT1 | Y | Y | N |
| *SYNE1* | ENSG00000284615: novel (AS) | Y | Y | N |
| *TGFBR2* | predicted (AS) | ENSMUSG00000132121 | Y | Y |
| *TNNT2* | ENSG00000286600: novel | Y | Y | Y |
| *TPM1* | ENSG00000259498: TPM1-AS | Y | Y | Y |
| *TTN* | ENSG00000270956: TTN-AS1 | ENSMUSG00000086354 | Y | Y |
| *VEGFA* | ENSG00000272114: novel (AS) | ENSMUSG00000128887 | Y | N |
| *XIRP1* | ENSG00000287620: novel (AS) | Y | ENSSSCG00000041387 | Y |
| *ADAMTS6* | ENSG00000294488: novel (AS) | Y | Y | N |
| *AKAP13* | ENSG00000259367: AKAP13-AS1 | Y | Y | N |
| *ANKRD17* | ENSG00000250220: ANKRD17-DT | Y | Y | N |
| *ATN1* | ENSG00000293728: novel (AS) | Y | Y | N |
| *ATP11A* | ENSG00000232684: ATP11A-AS1 | N | N | N |
| *ATP1B1* | ENSG00000237707: novel | Y | Y | N |
| *ATP2A2* | ENSG00000289311: novel (AS) | Y | Y | N |
| *BMPR2* | ENSG00000273456: novel (AS) | Y | Y | N |
| *BVES* | ENSG00000203808: BVES-AS1 | Y | Y | N |
| *CASZ1* | ENSG00000272078: novel (AS) | ENSMUSG00000086606 | Y | Y |
| *CD47* | ENSG00000279277: novel | Y | Y | N |
| *CDH5* | ENSG00000305055: novel (AS) | ENSMUSG00000110053 | Y | N |
| *COL15A1* | ENSG00000270412: COL15A1-AS1 | Y | Y | N |
| *COL1A1* | ENSG00000249406: TILAM | Y | Y | N |
| *COL1A2* | ENSG00000285090: novel (AS) | ENSMUSG00000122837 | ENSSSCG00000060665 | N |
| *CSRP2* | ENSG00000257910: novel (AS) | Y | Y | N |
| *DSTN* | ENSG00000277373: novel | N | N | N |
| *EGFL7* | ENSG00000228401: LNCEGFL7OS | Y | Y | N |
| *EIF4EBP1* | ENSG00000260949: novel | N | N | N |
| *F2R* | ENSG00000225407: novel (AS) | Y | Y | N |
| *F3* | ENSG0000028873: novel | Y | Y | N |
| *FBLN1* | ENSG00000238120: LINC01589 | ENSMUSG00000142768 | Y | N |
| *FBN2* | ENSG00000307925: novel (AS) | Y | Y | Y |
| *FBXW7* | ENSG00000270751: FBXW7-AS1 | Y | Y | Y |
| *FN1* | ENSG00000230695: FN1-DT | Y | Y | N |
| *FSTL1* | ENSG00000286735: novel (AS) | Y | Y | N |
| *FZD8* | ENSG00000287528: novel (AS) | Y | Y | Y |
| *GAS6* | ENSG00000233695: GAS6-AS1 | Y | Y | N |
| *GATA6* | GATA6-AS1: ENSG00000266010 | Gata6os: ENSMUSG00000097222 | ENSSSCG00000039279 | Y |
| *GJC1* | ENSG00000267505: novel | Y | Y | N |
| *GNAI2* | ENSG00000213600: novel (intronic) | Y | Y | N |
| *GNG5* | ENSG00000285851: novel *sense* | Y | Y | Y |
| *HINT1* | ENSG00000290020: novel | Y | Y | N |
| *HMGA1* | ENSG00000288879: novel | Y | Y | N |
| *HNRNPA1* | ENSG00000258344: novel | Y | Y | N |
| *HNRNPU* | ENSG00000273175: novel | Y | Y | Y |
| *IL6ST* | ENSG00000227908: IL6ST-DT | Y | Y | N |
| *KLF2* | ENSG00000267275: KLF2-DT | Y | Y | N |
| *LIMS1* | ENSG00000228763: LIMS1-AS1 | Y | Y | N |
| *MB* | ENSG00000268818: novel | N | N | N |
| *MCL1* | ENSG00000289457: novel | Y | Y | N |
| *MLLT10* | ENSG00000304565: novel AS | Y | Y | N |
| *MRPS12* | ENSG00000269486: ERVK9-11 | N | N | N |
| *MYH7* | ENSG00000258444: novel (AS) | Y | Y | Y |
| *NAV1* | ENSG00000310071: novel (AS) | Y | Y | N |
| *NDRG1* | ENSG00000289316: novel | Y | Y | N |
| *NDRG4* | ENSG00000294769: novel (AS) | Y | Y | N |
| *NDUFA6* | ENSG00000237037: NDUFA6-DT | Y | Y | N |
| *NFAT5* | ENSG00000274093: novel | Y | Y | N |
| *OBSCN* | ENSG00000162913: OBSCN-AS1 | Y | Y | Y |
| *PCNT* | ENSG00000289421: novel | N | N | N |
| *PHIP* | ENSG00000286340: novel (AS) | Y | Y | Y |
| *PKN2* | ENSG00000237505:PKN2-AS1 | Y | Y | Y |
| *PLA2G15* | ENSG00000260441: novel (AS) | Y | Y | Y |
| *POPDC2* | ENSG00000289134: novel | N | N | N |
| *PTPRM* | ENSG00000266767: novel (AS) | Y | Y | N |
| *QKI* | ENSG00000306330: novel (AS) | Y | Y | N |
| *RAMP2* | ENSG00000197291: RAMP2-AS1 | Y | Y | N |
| *RBM24* | ENSG00000287559: novel (AS) | Y | Y | Y |
| *RBMS1* | ENSG00000300196: novel (AS) | Y | Y | Y |
| *SETD5* | ENSG00000296784: novel | Y | Y | N |
| *SHC1* | ENSG00000271380: PYGO2 and SHC1 AS | Y | Y | N |
| *SLC2A4* | ENSG00000263342: novel (AS) | Y | Y | N |
| *SMAD5* | ENSG00000164621: SMAD5-AS1 | Y | Y | N |
| *SMARCD3* | ENSG00000272661: novel (AS) | Y | Y | N |
| *SMG1* | ENSG00000260352: SMG1-DT | Y | Y | N |
| *SOD1* | ENSG00000234509: SOD1-DT | N | N | N |
| *SORBS2* | ENSG00000233110: SORBS2-AS1 | ENSMUSG00000085440: Sorbs2os | Y | Y |
| *SOX4* | ENSG00000283480: novel (AS) | Y | Y | N |
| *SPEN* | ENSG00000179743: SPEN-AS1 | Y | N | N |
| *SPP1* | ENSG00000286618: novel (AS) | Y | Y | N |
| *SPTB* | ENSG00000288779: novel | Y | Y | N |
| *SRL* | ENSG00000262468: LINC01569 | Y | Y | N |
| *SYNM* | ENSG00000259475: SYNM-AS1 | Y | Y | N |
| *TMSB4X* | ENSG00000288817: novel | Y | Y | N |
| *TRIM63* | ENSG00000309383: novel (AS) | Y | Y | N |
| *UBR4* | ENSG00000272084: novel | Y | Y | N |
| *VCAN* | ENSG00000249835: VCAN-AS1 | Y | Y | Y |
| *VLDLR* | ENSG00000236404: VLDLR-AS1 | Y | Y | N |
| *ZFP36* | ENSG00000288919: novel | ENSMUSG00000120054 | Y | N |
| *ZFP36L1* | ENSG00000289626: novel | Gm56530 | Y | N |
| *ZMIZ1* | ENSG00000224596: ZMIZ1-AS1 | ENSMUSG00000087535 (Zmiz1os1) | ENSSSCG00000054028 | N |
| *ZNF280D* | ENSG00000260392: novel | Y | Y | N |

Abbreviations: Y, presence of microsynteny; N, absence of microsynteny
